# Genome-guided manipulation of regulators of morphogenesis in a *C. albicans* strain with low virulence is not sufficient to trigger a high-virulence phenotype

**DOI:** 10.1101/2025.07.16.665085

**Authors:** Ricardo Fróis-Martins, Sarah Mertens, Van Du T. Tran, Corinne Maufrais, Christophe d’Enfert, Dominique Sanglard, Salomé LeibundGut-Landmann

**Affiliations:** Section of Immunology, Vetsuisse Faculty, University of Zurich, Zurich, Switzerland; Institute of Experimental Immunology, University of Zurich, Zurich, Switzerland; Vital-IT, SIB Swiss Institute of Bioinformatics, Lausanne, Switzerland; Institut Pasteur, Université Paris Cité, INRAE USC2019, Unité Biologie et Pathogénicité Fongiques, Paris, France; Institute of Microbiology, University of Lausanne and University Hospital Center, Lausanne, Switzerland; Medical Research Council Centre for Medical Mycology at the University of Exeter, Department of Biosciences, Faculty of Health and Life Sciences, Exeter, UK

**Keywords:** *Candida albicans*, oral colonization, pathogenicity, virulence, hyphal network, cAMP-PKA pathway, *BRG1*

## Abstract

As a member of the microbiome, *C. albicans* colonizes the oral cavity and other mucosal surfaces of the human body. While commensalism is tightly controlled by the host immune system, the fungal determinants enabling the fungus to colonize the host mucosa without causing tissue damage and inflammation remain less clear. In search of genetic determinants that may underly the commensal lifestyle of the low-virulent *C. albicans* isolate 101, we identified a small sequence duplication in one of the *BRG1* allele resulting in a truncated loss-of-function allele (*BRG1^TRUNC^*). Replacing *BRG1^TRUNC^* by the full-length allele (*BRG1^FL^)* resulted in a modest increase in filamentation, but did not alter the phenotype of the fungus in the oral mucosa of experimentally colonized mice. Spontaneous outgrowth of highly virulent variants of the 101 strain with the *BRG1^TRUNC^* alleles replaced by a full-length allele identified the Cyr1-cAMP-PKA signalling pathway as a modulator of fungal virulence. Although the Glu-to-Lys mutation in *CYR1* (*CYR1^E1541K^*), which converts Cyr1 into a hyperactive adenylate cyclase, greatly increased filamentation, the expression of hyphae-associated genes, and host cell damage when tested *in vitro*, it was insufficient to increase virulence of *C. albicans* strain 101 in the oral mucosa *in vivo*, irrespective of the *BRG1* status. Together, this highlights that the low-virulent nature of strain 101 is firmly anchored and cannot be overcome by manipulating *BRG1* and *CYR1,* two genes with known roles in *C. albicans* virulence.

**IMPORTANCE:** During homeostasis, the fungus *Candida albicans* establishes mutualistic interactions with its human host. It can however also adopt a pathogenic state and cause infections with diverse clinical manifestions that pose a significant challenge for diagnosis and therapy. Understanding the fungal determinants that underlie *C. albicans* colonization under steady state conditions may thus provide new avenues for modulating the fungus-host interaction in candidiasis patients to restore homeostasis. Here, we report gene variants that distinguish high-virulent from low-virulent *C. albicans*. Gene-exchange mutants provided evidence for the contribution of a *BRG1* loss-of-function allele and a *CYR1* gain-of-function mutation toward fungal pathogenicity. However, *in vivo* in an experimental model of *C. albicans* oral colonization, none of these gene variants individually or in combination were sufficient to change the virulence profile of the fungus. These findings indicate that *C. albicans* mucosal colonization is regulated by a complex gene network rather than by single genetic determinants.

## INTRODUCTION

*Candida albicans* is a common member of the microbiome in the oral cavity, the gastrointestinal tract, and the female reproductive tract (1). Despite its primarily commensal lifestyle, *C. albicans* can become pathogenic and is therefore also referred to as a pathobiont. Under conditions of compromised host immunity or upon dysbiosis, it can cause superficial infections that manifest as oral and vaginal thrush (2, 3). In rare cases, such as in neutropenic individuals, *C. albicans* translocates across the epithelial barrier and disseminates via the bloodstream to cause systemic disease, which is associated with a high mortality rate due to liver, spleen and kidney failure, and/or fungal meningitis (2). The pathogenic impact of *C. albicans* further extends to non-infectious diseases such as inflammatory bowel disease and ethanol-induced liver disease (4–8). Treatment options are limited and the rise in antifungal resistance jeopardises the effectiveness of the few available antifungal drugs (9). A better understanding of the mechanisms governing fungal pathogenicity is needed to guide new approaches for disease prevention and therapy by targetting fungal pathogenicity determinants rather than aiming at eradicating the oragnisms with fungicidal drugs.

A hallmark of *C. albicans* pathogenicity is its capacity to undergo morphological changes and switch between yeast and hyphal forms. While originally, the yeast form was primarily associated with colonization and the hyphal morphotype with pathogenicity, this dichotomy has been challenged by the observation that both, yeast-locked and hyperfilamentous mutants are avirulent in experimental infection models (10–12). Moreover, we and others have recently found that homeostatic colonization of the oro-gastrointestinal tract also depends on the capacity of *C. albicans* to filament, while yeast-locked mutants are bad colonizers (10, 12, 13). Therefore, dynamic morphological changes are critical for both, fungal colonization and disease. Limiting the degree of filamentation and the processes associated with filamentation is essential for stable colonization during steady state and for tissue homeostasis (13, 14)

1. *C. albicans* filamentation is regulated by environmental cues that are integrated by diverse signal transduction pathways, including the MAPK pathway, the Rim101 pathway, and the cAMP-dependent potein kinase A (PKA) pathway (15, 16). The adenylate cyclase Cyr1 plays a central role in translating environmental cues into PKA activation, in part by Ras-dependendent mechanisms (17). Cyr1 converts ATP into the secondary messenger cAMP, which leads to PKA activation and in turn phosphorylation of the master regulator of the yeast-to-hyphae transition, Efg1 (18). Activated Efg1 not only drives the core filamentation response in *C. albicans* but also induces expression of numerous virulence genes, including *HWP1, ALS3*, and *ECE1*. These hyphae-associated genes mediate fungal adhesion, tissue invasion, and host cell lysis, key processes in the interaction of the fungus with the host culminating in infection and tissue damage (19–21).

To limit the expression of these virulence traits*, C. albicans* possesses hyphal repressors such as Nrg1, a DNA-binding protein repressing filamentous growth via the Tup1 pathway (22, 23). The expression of *NRG1* itself is regulated by Brg1, which influences the stability of *NRG1* transcripts, thus controlling hyphae formation and virulence via a feedback circuit (11).

*C. albicans* exhibits a large intraspecies diversity (24) with strain-specific differences at the genomic, genetic, and epigenetic levels translating into phenotypic variations (25, 26). As such, the *C. albicans* species comprises a large spectrum of isolates differing in their intrinsic degree of virulence, which in turn is further modulated by environmental and host factors. Most experimental studies have been conducted with strain SC5314, although this strain is not fully representative of the species due to its intrinsic high virulent properties (25, 26). When experimentally administered to mice, it causes an acute inflammatory response and is rapidly cleared (27). More recently, different groups including our own started to explore the nature of low-virulent strains and their behavior at the host interface (25, 28, 29). Pinpointing the molecular basis underlying the phenotypic differences between strains is challenging given the number of genetic differences that often separate natural isolates of *C. albicans* (30).

The oral isolate 101 is a low-virulent strain suitable for studying long-term fungal mucosal colonization. It persistently colonizes the oral mucosa of immunocompetent and non-antibiotically treated mice without causing tissue damage and inflammation, reminiscent of *C. albicans* homeostatic colonization in humans. Yet, T cell and IL-17 deficiency causes overgrowth of isolate 101 leading to symptoms reminiscent of oral thrush (25, 31, 32). The intrinsically low-virulent nature of the strain is explained by the strong attenuation of key virulence genes, including those linked to filamentation owing to high expression of the transcriptional repressor gene *NRG1* (14). Deletion or repression of *NRG1* rendered strain 101 more virulent by derepressing the hyphal program (14). In search of genetic determinants underlying the elevated expression of *NRG1* in strain 101, we identified a truncated *BRG1* allele that was not present in other low-virulent *C. albicans* isolates. We defined the role of the truncated allele in fungal pathogenicity *in vitro* and *in vivo*. Furthermore, we found that strain 101 spontaneously acquires mutations in the RAS/cAMP-PKA pathway, which derepressed several virulent traits. The strong induction of filamentation, *ECE1* expression and epithelial damage observed under *in vitro* conditions was however not paralleled by increased pathogenicity *in vivo*. Instead, strain 101 variants preserved a largely low-virulent behavior in the murine oral mucosa, highlighting that the low-virulent nature of strain 101 is firmly anchored and cannot be overcome by manipulating individual genes associated with virulence.

## RESULTS

### The truncated *BRG1* allele identified in the low-virulent strain 101 is unable to drive virulence of strain SC5314

In search of mutations that may underly the low-virulent phenotype of strain 101, we established and annotated the full genome of strain 101 (Bioproject PRJNA923600). The 101 genome was resolved into 2 separate haplotypes (Bioprojects PRJNA1074522 and PRJNA1074523; see Material and Methods for details) with each 8 large contigs corresponding to the 8 chromosomes of the reference SC5314. The 2 haplotypes contained 5829 and 5816 annotated genes, which is lower than SC5314 (6213 annotated genes) but principally due to the absence of annotations for ORF smaller than 500 bp in the 101 haplotypes. The two 101 haplotypes exhibit low homozygosity (approximately 11’000 SNPs between each haplotype).

While focusing on known *C. albicans* genes affecting virulence and/or filamentation, we identified using haplotype alignment a 209 base pair duplication at the 3’ end of the *BRG1* gene in one of the haplotypes, creating a truncated protein due to a frameshift (**Figure 1A**). Amplification by PCR of *BRG1* alleles in strain 101 confirmed the presence of two distinct alleles that differ in length (**Figure 1B**). The gene duplication responsible for the truncated *BRG1* allele in strain 101 (*BRG1*^TRUNC^) is unique across >250 whole-genome (short-read) sequenced isolates.

**Figure 1.**
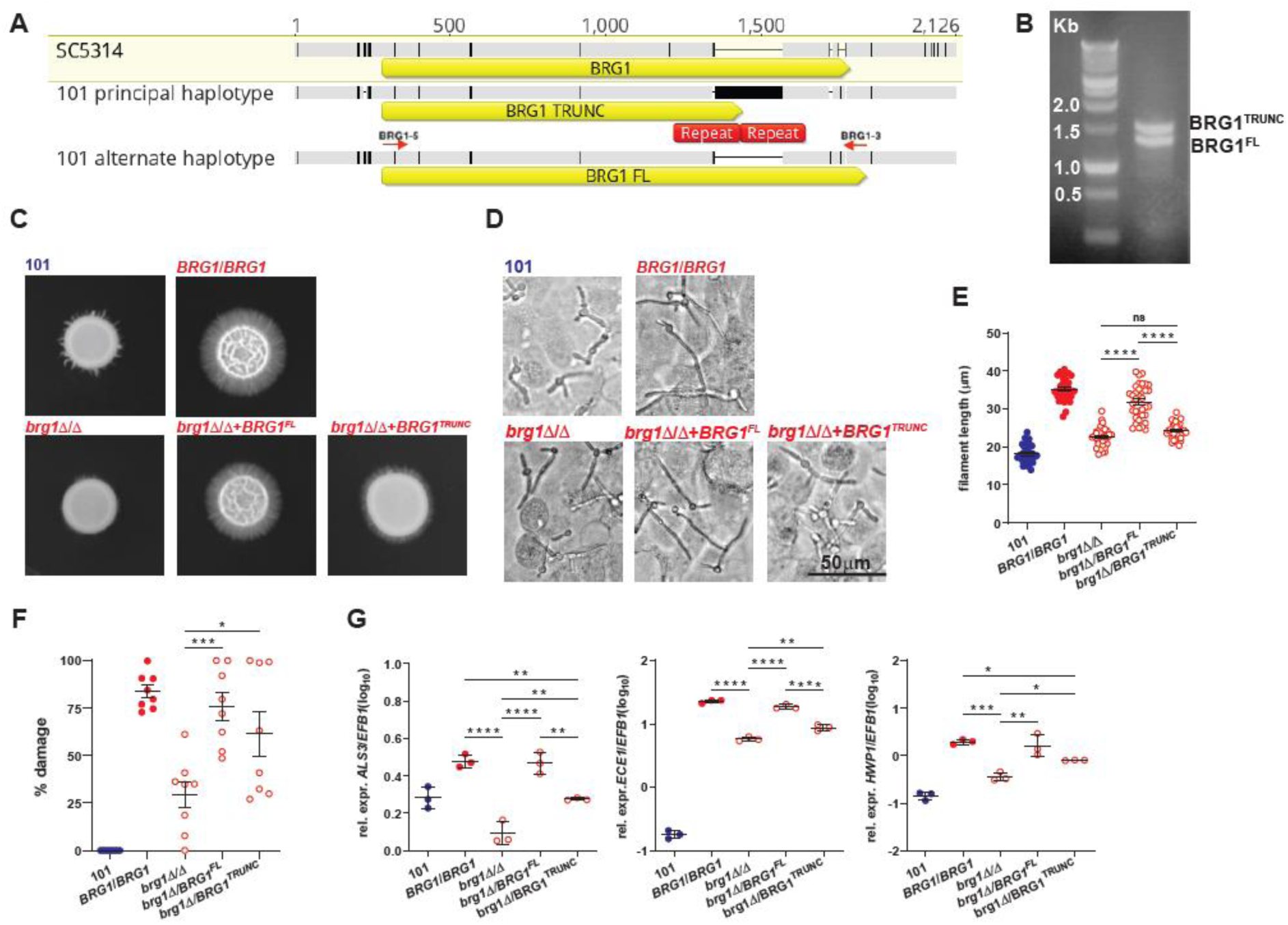
The truncated *BRG1* allele identified in the low-virulent strain 101 is unable to drive virulence of strain SC5314. **A.-B.** Two different *BRG1* alleles are present in strain 101. **A.** Alignment of the *BRG1* region of the two haplotypes of strain 101 compared to the *BRG1* locus (C1_05140W_A) of SC5314 as reference (yellow highlighted)).. *BRG1* open reading frames (ORF) are indicated by yellow arrows below each aligned nucleotide sequence. SC5314 BRG1 region The polymorphisms between the different *BRG1* alleles are indicated by black bars in the greyed nucleotide sequences The repeated regions (indicated by red rectangles) results in a 200-bp increase of PCR product for allele BRG1-TRUNC (see planel B).Alignment was obtained with Geneious Prime (Biomatters, Ltd.). **B.** PCR amplification of *BRG1* alleles in 101 and visualization of two different band sizes (1.5 and 1.3 kb). Primers BRG1-5 and BRG1-3 that were used in PCR are showed in Panel A and are located at the boundaries of the *BRG1* orf, resulting in 1.5 and 1.3 kb products for BRG1-TRUNC and BRG1-FL, respectively, which is consistent with gel electrophoresis. Left lane: molecular weight standard (Benchtop 1 kb DNA Ladder, Promega). **C.-G.** *C. albicans* strains SC5314_*BRG1*/*BRG1*, SC5314_*brg1*Δ/Δ, SC5314_*brg1*Δ/*BRG1*^FL^, SC5314_ *brg1*Δ/*BRG1*^TRUNC^, and 101 were assessed for their phenotype *in vitro*. **C.** Colony morphology of the strains grown on Spider agar for 7 days at 35°C**. D.** Representative images of filamenting strains upon contact with a monolayer of TR146 keratinocytes for 3.5 hours at 37°C, 5% CO_2_. **E.** Quantification of the filament length. Each symbol represents the mean filament length of 20 – 30 filaments per field. Data are pooled from two independent experiments with 15-20 visual fields analyzed per strain each. The mean ± SEM is indicated. **F.** LDH released from TR146 keratinocytes after exposure to the fungal strains for 24 hours at 37°C, 5% CO_2_. Each symbol represents one well. Data are pooled from two independent experiments with 4 wells per strain each. The mean ± SEM is indicated. **G.** *ECE1*, *HWP1*, and *ALS3* expression levels by the fungal strains after exposure to TR146 keratinocytes for 24 hours at 37°C, 5% CO_2_. Each symbol represents one sample. Data are from one representative out of two independent experiments with three samples per strain each. The mean ± SD is indicated. Statistical significance was determined using one-way ANOVA. *p<0.05, **p<0.01, ***p<0.001, ****p<0.0001. **See also Figure S1**.

The second haplotype retained a full length copy of the *BRG1* gene in strain 101 (termed *BRG1-FL* in the following), which slightly differs from the two revised *BRG1* alleles of strain SC5314 (**Suppl. Figure S1**). Notably, our phasing of single nucleotide polymorphisms and analysis of insertion-deletion data at the *BRG1* locus for 182 *C. albicans* genome-sequenced isolates (Ropars et al., 2018) revealed a frameshift in the *BRG1* haplotypes of strain SC5314 (Assembly 22). Correction of this frameshift as well as glutamine-rich coding regions led to two Brg1 proteins with C-terminal sequences identical to that in the Brg1-FL protein of strain 101 (**Suppl. Figure S1**). The same was observed for strains CEC3672 and CEC3678 that will be described below. Given the role of Brg1 as a transcription factor associated with *C. albicans* virulence and biofilm formation acting as negative regulator of Nrg1 (11, 33) and because of the causal link between the degree of *NRG1* expression and the low-virulent phenotype of strain 101 (14), we decided to experimentally investigate the functional role of the newly identified truncated *BRG1* allele in *C. albicans* pathogenicity.

We started by introducing the truncated allele, or the full length allele from strain 101 (*BRG1*^FL^) as a control, into a *BRG1* null mutant of strain SC5314 (SC5314_*brg1*ΔIΔ). *BRG1* expression levels were restored irrespective of the allele (**Suppl. Figure S2A**). While full deletion of *BRG1* strongly impacted filamentation, introduction of one copy of the full-length *BRG1* allele of strain 101 into SC5314_*brg1*Δ*I*Δ was sufficient to restore filamentation on Spider agar (**Figure 1C**), indicating that *BRG1*^FL^ of strain 101 is fully functional. In contrast, the truncated *BRG1* allele of strain 101 was unable to do so (**Figure 1C**). Likewise, the filamentation defect of strain SC5314_*brg1*Δ*I*Δ when placed in contact with keratinocytes in FCS-containing medium was restored upon introduction of *BRG1*^FL^ in this strain while it was not restored upon introduction of *BRG1*^TRUNC^ (**Figure 1D-E**). Introduction of the *BRG1*^TRUNC^ allele into wild-type SC5314, replacing one full-length allele while leaving intact the second full-length allele, did not compromise filamentation under the same assay conditions, indicating that the *BRG1*^TRUNC^ allele did not act as a dominant-negative allele (**Suppl. Figure S2B**).

In line with the requirement of strong filamentation for epithelial cell damage induction by *C. albicans* SC5314 (19), introducing the *BRG1*^TRUNC^ allele into SC5314_*brg1*Δ/Δ partially restored the high degree of damage in TR146 keratinocytes that was elicited in response to strains containing one or two copies of the full-length allele (**Figure 1F**). Again, in a mutant strain containing both, a *BRG1*^TRUNC^ and a full length *BRG1* allele, the truncated allele did not alter the effect of the full-length allele (**Suppl. Figure S2C**).

The dependence of the virulence phenotype on *BRG1* integrity extended to the expression of fungal virulence genes, including the candidalysin-encoding *ECE1* gene as well as the adhesin-coding genes *ALS3* and *HWP1*, all among the genes that were defined to constitute the core filamentation response (34). While the introduction of *BRG1*^FL^ into SC5314_*brg1*ΔIΔ restored expression levels similar to those in wild-type SC5314, the *BRG1*^TRUNC^ allele led to only limited expression of hyphae-associated genes (**Figure 1G**). In presence of a full length gene copy, introduction of *BRG1*^TRUNC^ had not impact on virulence gene expression, again confirming that he truncated allele was not dominant-negative (**Suppl. Figure S2D**). Together, these results demonstrate that the truncated *BRG1* allele identified in the low-virulent strain 101 was dysfunctional in *C. albicans*. Although our experiments demonstrate that it does not act as a dominant negative allele in the high-virulent strain SC5314, we speculate that in a low-virulent strain such as strain 101, it may impact the strain’s virulence, even in presence of an intact full-length allele.

### Restoring the full-length *BRG1* allele in the low-virulent strain 101 is insufficient to increase the strain’s virulence

After having found that the truncated *BRG1* allele of strain 101 is hypomorphic, while the full-length allele is fully functional in the context of the highly virulent strain SC5314 (**Figure 1**), we speculated that replacing the truncated allele in strain 101 with a second copy of the full length allele may induce virulence traits in this low-virulent strain. Notably, a strain 101 derivative bearing two *BRG1*^FL^ alleles did not show any change in colony morphology on Spider agar, when compared to the parental strain 101 bearing a *BRG1*^TRUNC^ allele and a *BRG1*^FL^ allele (**Figure 2A, top and Suppl. Fig. S3**). Likewise, the replacement of the *BRG1*^TRUNC^ allele by the *BRG1*^FL^ allele did not increase the degree of filamentation of strain 101 when placed in contact to TR146 keratinocytes in FCS-containing medium (**Figure 2B, C**). Growth on YCB/BSA agar, which also triggers filamentation (embedded growth), indicated that the allele replacement did however have a small impact on fungal morphogenesis (**Figure 2A, bottom**). Moreover, longer incubation on Spider agar at 37°C also revealed the mildly enhanced filamenting capacity of the mutant (**Figure 2D**). However, the effect was not paralleled by a restoration of virulence. Epithelial cell damage induction, which serves as a good correlate for *C. albicans* pathogenicity in barrier tissues (25), was not enhanced by replacement of the *BRG1*^TRUNC^ allele by the *BRG1*^FL^ allele in strain 101 (**Figure 2E**). Likewise, expression of the core virulence genes *ECE1*, *HWP1*, and *ALS3* was not enhanced and *NRG1* levels remained highly expressed (**Figure 2F**). Together, these findings indicate that replacing the hypomorphic *BRG1*^TRUNC^ allele in strain 101 by a fully functional copy of *BRG1* was not sufficient to alter the strain’s pathogenicity with respect to damage induction and virulence gene expression and only exerted a very limited effect on morphology when grown under *in vitro* conditions.

**Figure 2.**
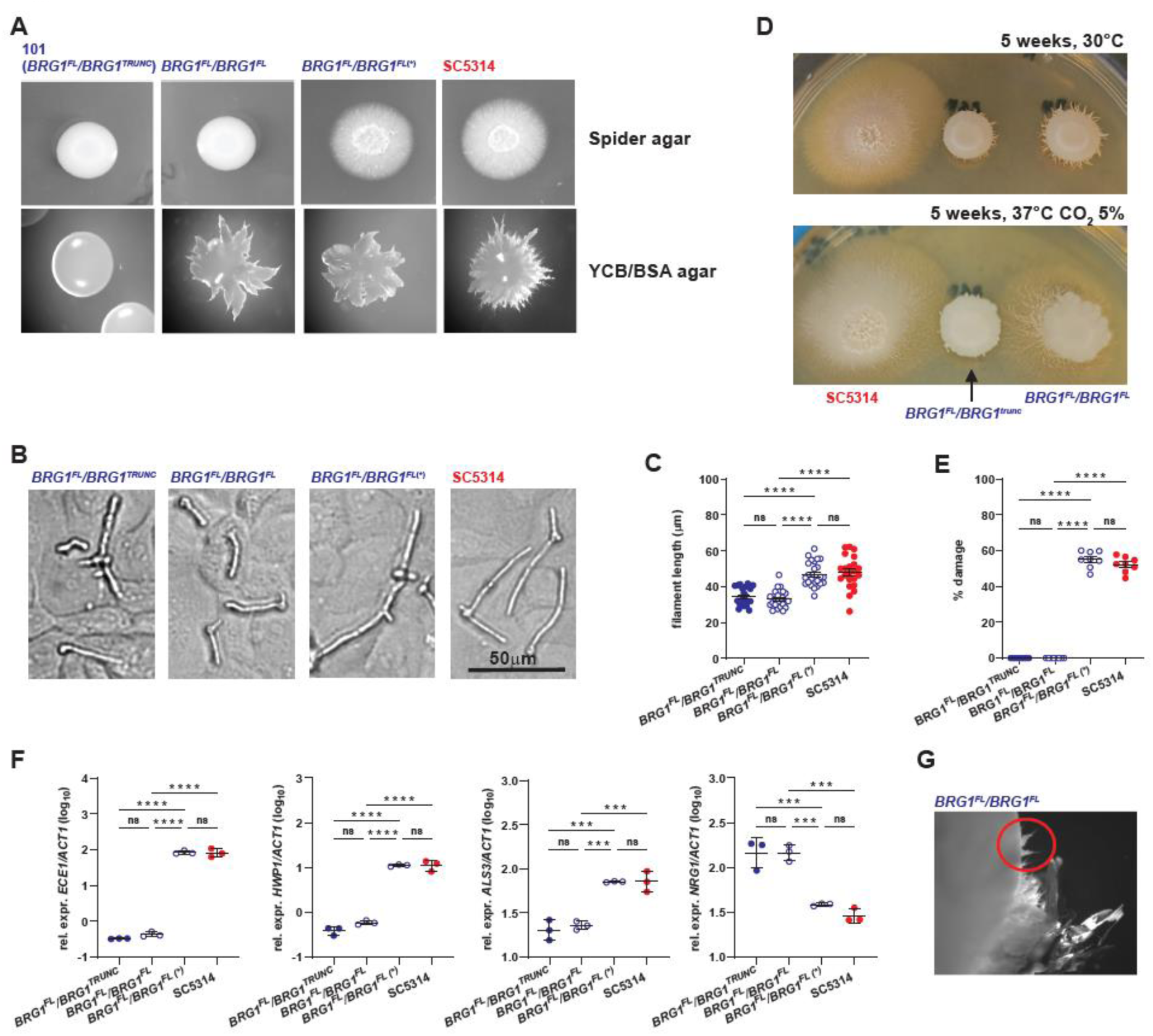
Restoring the full-length *BRG1* allele in low-virulent strain 101 is insufficient to increase the strain’s virulence when tested *in vitro*. *C. albicans* strains 101_*BRG1*^FL^/*BRG1*^TRUNC^, 101_*BRG1*^FL^/*BRG1*^FL^, 101_*BRG1*^FL^/*BRG1*^FL^(*), and SC5314 wild type were assessed for their phenotype *in vitro*. **A.** Colony morphology of the strains grown on Spider agar (top) or on YCB/BSA agar (bottom) for 7 days at 35°C**. B.** Representative images of filamenting strains upon contact with a monolayer of TR146 keratinocytes for 3.5 hours at 37°C, 5% CO_2_. **C.** Quantification of the filament length. Each symbol represents the mean filament length of 20 – 30 filaments per visual field. Data are pooled from two independent experiments with 10-12 visual fields analyzed per strain each. The mean ± SEM is indicated. **D.** Colony morphology of the indicated strains grown on Spider agar for 5 weeks at 37°C, 5% CO_2_**. E.** LDH release from TR146 keratinocytes after exposure to the fungal strains for 24 hours at 37°C, 5% CO_2_. Each symbol represents one well. Data are pooled from two independent experiments with 4 wells per strain each. The mean ± SEM is indicated. F. *ECE1*, *HWP1*, *ALS3*, and *NRG1* expression levels by the fungal strains after exposure to TR146 keratinocytes for 24 hours at 37°C, 5% CO_2_. Each symbol represents one sample. Data are from one representative out of two independent experiments with three samples per strain each. The mean ± SD is indicated. **G.** Close-up of a colony of 101_*BRG1*^FL^/*BRG1*^FL^ on Spider agar showing signs of filamentation in a subcolony. In C, D, F and G, statistical significance was determined using one-way ANOVA. *p<0.05, **p<0.01, ***p<0.001, ****p<0.0001. **See also Figure S2**.

### Spontaneous mutants convert the low-virulent strain 101 in a highly virulent variant

While performing *in vitro* experiments with strains 101_*BRG1*^FL^/*BRG1*^TRUNC^ and 101_*BRG1*^FL^/*BRG1*^FL^ shown in **Figure 2A-F**, we observed that the latter strain, when grown on Spider agar, displayed some spikes at the border of the colony, resembling hyphal outgrowth (**Figure 2G**). Material from the spike was collected and diluted to obtain single colonies. This strain, termed 101_*BRG1*^FL^/*BRG1*^FL^(*) displayed a highly filamentous morphology under all conditions tested. On Spider agar, in suspension and in contact to keratinocytes, the strain closely resembled the high virulent strain SC5314 (**Figure 2A-C and Suppl. Fig. S3**). It also elicited a strong LDH release response in TR146 cells (**Figure 2E**), and it displayed a gene expression profile confusingly similar to that of strain SC5314 (**Figure 2F**). Together, these findings indicate that a low-virulent strain can convert into a highly virulent variant as inferred from *in vitro* conditions.

To further investigate the pathogenicity of the spontaneously outgrown 101_*BRG1*^FL^/*BRG1*^FL^(*) strain at the interface with the host, we made use of our experimental model of oral colonization in mice. In this model, *C. albicans* efficiently establishes colonization of the epithelium, as can be assessed by quantification of the fungal burden in the tongue and by histological analysis of the tongue epithelium (25). Highly virulent strains such as SC5314 swiftly induce a strong inflammatory response upon invasion of fungal hyphae into the nucleated layers of the epithelium (stratum granulosum, stratum spinosum) where they stimulate cytokine and chemokine production, which in turn results in massive accumulations of inflammatory cells. This massive response, which peaks on day 1-2 post-infection, leads to clearance of the fungus within 3-5 days (27, 35, 36). In contrast, low virulence strains such as 101 reside predominantly in the stratum corneum and elicit a non-inflammatory homeostatic response that is characterized by tissue-resident Th17 cells (13, 25). *C. albicans*-specific Th17 cells mediate immunosurveillance to prevent fungal overgrowth, without eliminating the fungus, thereby allowing long-lasting colonization, which mimics *C. albicans* colonization in humans. This model is therefore ideally suited for assessing and distinguishing high and low pathogenic isolates of *C. albicans* (25).

We colonized wild type C57BL/6 mice with strain 101 carrying one or two *BRG1*^FL^ alleles as well as the spontaneously outgrown 101_*BRG1*^FL^/*BRG1*^FL^(*) strain in comparison to SC5314. All strains colonized the tongue with comparable efficiency (**Figure 3A**). Reminiscent of what we observed *in vitro*, virulence factor genes were strongly induced and *NRG1* expression repressed in 101_*BRG1*^FL^/*BRG1*^FL^(*), but not in 101_*BRG1*^FL^/*BRG1*^TRUNC^ or 101_*BRG1*^FL^/*BRG1*^FL^ (**Figure 3B**). Strong induction of virulence factor gene expression correlated with the activation of a pronounced inflammatory response. Large agglomerates of inflammatory cells accumulated in the tongue epithelium of mice colonized with high-virulent isolates within as little as 1 day after infection (**Figure 3C**). Concomittantly, we observed expression of inflammatory cytokines in the colonized tissue (**Figure 3D**). Again, this response was limited to the spontaneously arisen high virulent mutant and its extent was comparable to that of the well-charachterized strain SC5314. Reminiscent of what is known for other high-virulent strains including SC5314, strain 101_*BRG1*^FL^/*BRG1*^FL^(*) did not persist, but was rapidly cleared from the tongue epithelium, despite its origin from the persistent colonizer 101 (**Figure 3E**). Together, this indicates that spontaneous changes converted a low-virulent strain into a high virulent one that phenocopies the prototypical high-virulent strain SC5314 in all parameters tested.

**Figure 3.**
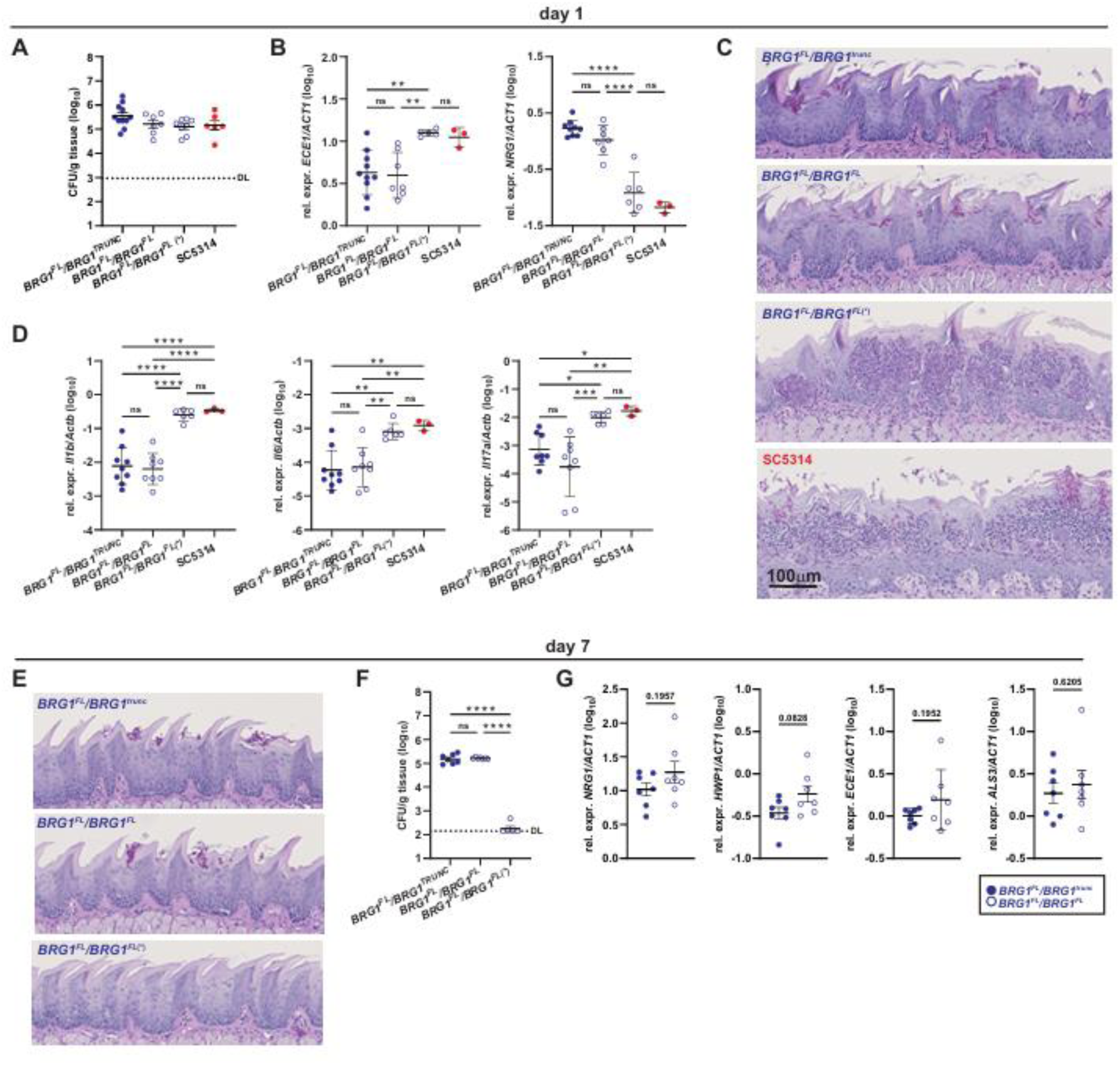
**Restoring the full-length *BRG1* allele in low-virulent strain 101 is insufficient to increase the strain’s virulence *in vivo*.**C57BL/6 mice were colonized with *C. albicans* strains 101_*BRG1*^FL^/*BRG1*^TRUNC^, 101_*BRG1*^FL^/*BRG1*^FL^, 101_*BRG1*^FL^/*BRG1*^FL^(*), and SC5314 wild type for 1 day (A-D) or 7 days (E-G). **A.** Tongue fungal burden on day 1 after colonization. DL, detection limit. **B.** *ECE1* and *NRG1* expression levels in the tongue tissue on day 1 after colonization. **C.** PAS-stained histology sections of the tongue on day 1 after colonization. **D.** Cytokine expression levels in the tongue tissue on day 1 after colonization. **E.** Tongue fungal burden on day 7 after colonization. DL, detection limit. **F.** PAS-stained histology sections of the tongue on day 7 after colonization. **G.** *ECE1*, *ALS3*, *HWP1* and *NRG1* expression levels in the tongue tissue on day 7 after colonization. In A, B, D, E, G, and H, each symbol represents one animal. Data are pooled from at least two independent experiments except for SC5314 in B and D. The mean ± SEM is indicated. Statistical significance was determined using two-way ANOVA. (B, D, E) or unpaired Student’s *t* test (G). *p<0.05, **p<0.01, ***p<0.001, ****p<0.0001. **See also Figure S3**.

The mutant of 101 carrying two full-lenth *BRG1* alleles could not be distinguished from its parental carrying only one fully functional *BRG1* allele in terms of its behavior in the murine tongue tissue based on fungal burden, histology, and expression of fungal virulence genes and host genes on day 1 post-infection (**Figure 3A-D**). Accordingly, the strain persisted in the tongue as did the parental 101_*BRG1*^FL^/*BRG1*^TRUNC^ with high fungal burden one week after infection (**Figure 3E, F**). Of note, no significant differences were detected in the expression of *ECE1*, *HWP1*, and *ALS3* in the tongue colonized with strain 101_*BRG1*^FL^/*BRG1*^FL^ in comparison to the parental 101_*BRG1*^FL^/*BRG1*^TRUNC^ on day 7 post-infection (**Figure 3G**).

### *CYR1*^E1541K^ induces the expression of virulence traits in the low-virulent strain 101

We hypothesized that the high virulence of strain 101_*BRG1*^FL^/*BRG1*^FL^(*) was the result of spontaneous mutations arising during growth of strain 101_*BRG1*^FL^/*BRG1*^FL^ on Spider medium. To identify such mutation(s), whole genome sequencing of 101_*BRG1*^FL^/*BRG1*^FL^(*) was performed and the genome was compared to that of the initial parent strain, 101_*BRG1*^FL^/*BRG1*^FL^. Among the detected changes (**Suppl. Figure S4 and Suppl. Table S1**), we noticed a non-synonymous point mutation in the gene encoding the adenlyl cyclase Cyr1 that leads to a Glu to Lys substitution in a conserved C-terminal motif close to the protein’s catalytic domain (position 1541). As an integral part of the cAMP-PKA pathway, Cyr1 regulates *C. albicans* morphogenesis (18). In response to environmental cues, it generates short-lived intracellular cAMP spikes. The Glu to Lys substitution at position 1541 of Cyr1 has been described previously to render the enzyme hyperactive and to cause constitutive filamentous growth of *C. albicans* even under non-hyphae-inducing conditions (37).

To assess whether the *CYR1*^E1541K^ point mutation was indeed responsible of the high-virulence phenotype of the spontaneously outgrown strain 101_*BRG1*^FL^/*BRG1*^FL^(*), we used CRISPR/Cas9 to introduce the mutation directly into the wild type 101 strain (**Suppl. Figure S4**). Modification of one allele was sufficient to drive a strong filamentation phenotype on Spider agar (**Figure 4A**) and in contact with TR146 keratinocytes (**Figure 4B, Suppl. Figure S5A**). Increased filamentation was accompanied by repression of *NRG1* and strong induction of hyphae associated genes to levels approximating or even exceeding those measured in SC5314 (**Figure 4C, Suppl. Figure S5B**). Despite strong filamentation and high levels of *ECE1* expression, the 101_*CYR1*^E1541K^ mutant did not phenocopy strain SC5314 regarding epithelial cell damage induction(**Figure 4D**).

**Figure 4.**
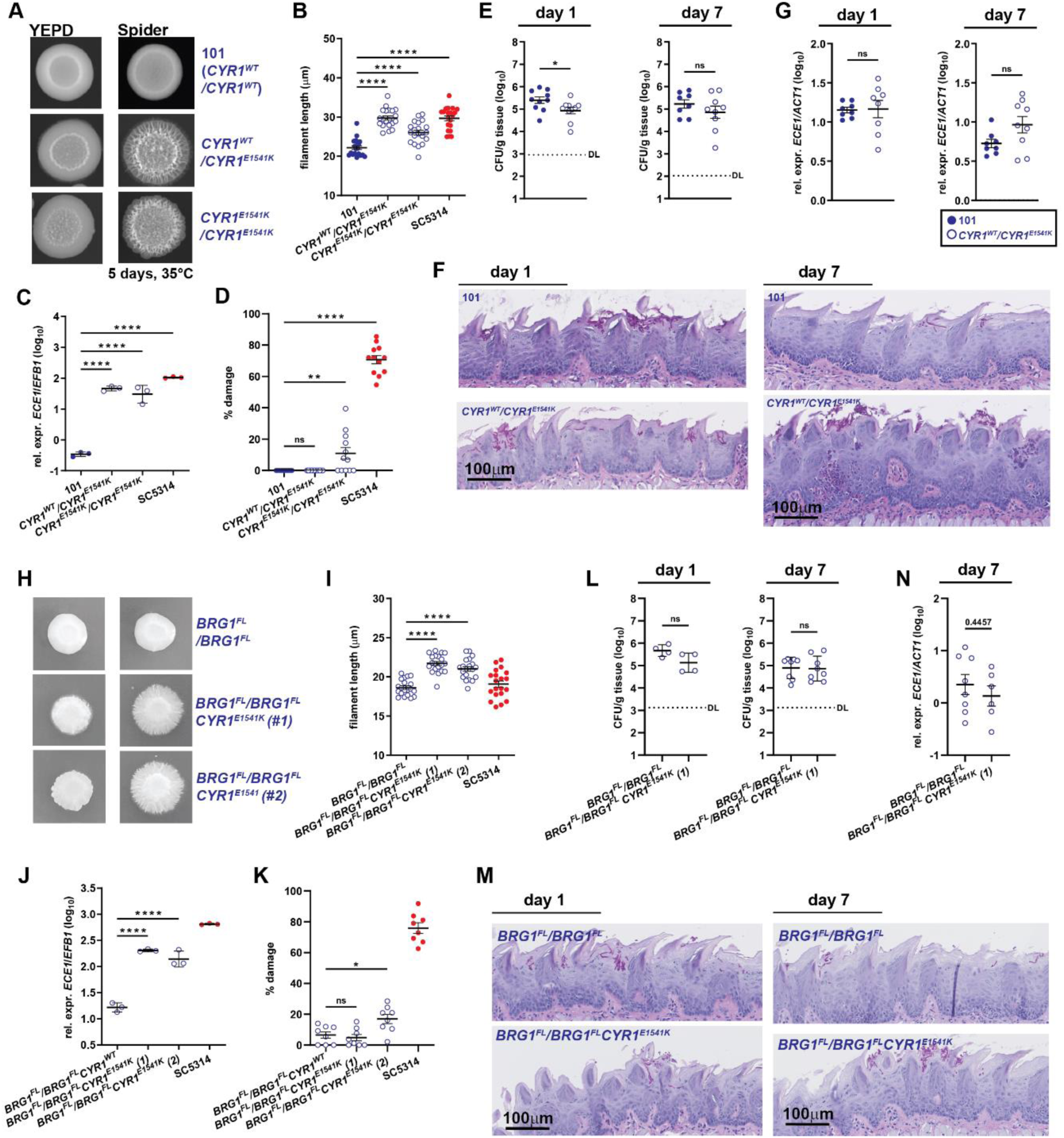
***CYR1*^E1541K^ induces the expression of virulence traits in the low virulent strain 101. A.-D.** *C. albicans* strains 101 wild type, 101_*CYR1*^E1541K^, and 101_*CYR1*^E1541K/E1541K^ were assessed for their phenotype *in vitro*. **A.** Colony morphology of the strains grown on Spider agar for 5 days at 35°C**. B.** Filamentation of the strains upon contact with a monolayer of TR146 keratinocytes for 3.5 hours at 37°C, 5% CO_2_. Each symbol represents the mean filament length of 20 – 30 filaments per visual field. Each symbol represents the mean filament length of 20 – 30 filaments per visual field. The mean ± SEM is indicated. **C.** *ECE1* expression levels by the fungal strains after exposure to TR146 keratinocytes for 24 hours at 37°C, 5% CO_2_. Each symbol represents one sample. Data are from one representative out of two independent experiments with three samples per strain each. The mean ± SD is indicated. **D.** LDH release from TR146 keratinocytes after exposure to the fungal strains for 24 hours at 37°C, 5% CO_2_. Each symbol represents one well. Data are pooled from two independent experiments. The mean ± SEM is indicated. **E.-G.** C57BL/6 mice were colonized with *C. albicans* strains 101 wild type or 101_*CYR1*^E1541K^ and tongue fungal burden (E), PAS-stained histology sections of the tongue (F), and *ECE1* expression levels in the tongue tissue were analyzed on day 1 and day 7 after colonization. Each symbol represents one mouse. Data are pooled from two independent experiments. The mean ± SEM is indicated. DL, detection limit. **H.-K.** *C. albicans* strains 101_*BRG1*^FL/^*BRG1*^FL^, 101_*BRG1*^FL/^*BRG1*^FL^ *CYR1*^E1541K^ (clone #1 and clone #2), and SC5314 wild type were assessed for their phenotype *in vitro*. **H.** Colony morphology of the strains grown on Spider agar for 5 days at 37 °C**. I.** Filamentation of the strains put in contact with a monolayer of TR146 keratinocytes for 3.5 hours at 37°C, 5% CO_2_. Each symbol represents the mean filament length of 20 – 30 filaments per visual field. The mean ± SEM is indicated. **J.** *ECE1* expression levels by the fungal strains after exposure to TR146 keratinocytes for 24 hours at 37°C, 5% CO_2_. Each symbol represents one sample. Data are from one representative out of two independent experiments with three samples per strain each. The mean ± SD is indicated. **K.** LDH release from TR146 keratinocytes after exposure to the fungal strains for 24 hours at 37°C, 5% CO_2_. Each symbol represents one well. . **L.-N.** C57BL/6 mice were colonized with *C. albicans* strains 101_*BRG1*^FL/FL^ and 101_*BRG1*^FL/FL^ *CYR1*^E1541K^ (clone 1). **L.** Tongue fungal burden on day 1 and day 7 after colonization. DL, detection limit. **F.** PAS-stained histology sections of the tongue on day 1 and day 7 after colonization. **G.** *ECE1* expression levels in the tongue tissue on day 7 after colonization. Each symbol represents one mouse. Data are pooled from two independent experiments. The mean ± SEM is indicated. Statistical significance was determined using one-way ANOVA (B-D and I-K) or unpaired Student’s *t* test (E-E, G, L-N). *p<0.05, **p<0.01, ***p<0.001, ****p<0.0001. **See also Figure S4 and Figure S5.**

The lack of high damage induction *in vitro* correlated with the inability of the 101_*CYR1*^E1541K^ mutant to drive inflammation *in vivo* in the oral mucosa of experimentally colonized mice (**Figure 4E-F**), as it is typical for highly virulent strains such as SC5314 (**Figure 3A-D**). The morphology of the 101_*CYR1*^E1541K^ mutant in the murine tongue tissue was indistinguishable from that of the parental 101 wild type strain, although quantification of filament length is impossible on tissue sections (**Figure 4F**). Likewise, the expression of hyphae-associated genes by the *CYR1*^E1541K^ mutant in comparison to the parental 101 wild type strain was not enhanced in the tongue tissue (**Figure 4G**). We were unable to test the homozygous *CYR1*^E1541K^ mutant *in vivo* because this strain formed aggregates as a result of hyperfilamentous phenotype, which precluded reproducible colonization of the murine oral mucosa. By day 7, when high virulent strains such as SC5314 are cleared from the experimentally infected tongue (25), fungal counts of the 101_*CYR1*^E1541K^ mutant were still high and indistinguishable from those of the parental wild type strain (**Figure 4E**). Fungal elements exhibited a mixed morphology. In very rare occasions we found larger fungal accumulations at the tongue surface, which in some instances also conincided with small inflammatory foci in the epithelium, whereby this was only the case for the *CYR1*^E1541K^ bearing mutant but not the wild type 101 strain (**Figure 4F**). This observation coincided with a small trend towards higher *ECE1* and reduced *NRG1* expression in the tongue tissue (**Figure 4G and Suppl. Fig 5C**).

The lack of virulence of the 101_*CYR1*^E1541K^ strain *in vivo* led us to question wether the full virulence-inducing potential of the hyperactive *CYR1*^E1541K^ gene variant was dependent on the presence of two intact *BRG1* alleles, as the high-virulent *CYR1*^E1541K^-carrying mutant originally emerged from the 101_*BRG1*^FL^/*BRG1*^FL^ strain (**Figure 2G**). We therefore introduced the *CYR1*^E1541K^ mutation into this strain to obtain a reconstituted mutant. This resulted in a strain that largely phenocopied 101_*CYR1*^E1541K^. It exhibited strongly enhanced filamentation in both Spider agar and in contact with TR146 keratinocytes and it strongly expressed hypae-induced genes (**Figure 4H-J, Suppl. Figure S6A**). However, these changes did not translate in enhanced keratinocyte damage induction if compared to the parental strain 101_*BRG1*^FL^/*BRG1*^FL^ (**Figure 5K**). Likewise, in the oral mucosa of experimentally colonized mice, the *CYR1*^E1541K^ variant did not exert a stronger virulence-inducing effect when introduced into 101_*BRG1*^FL^/*BRG1*^FL^ than in the wild type 101 background (**Figure 5L-N**).

**Figure 5.**
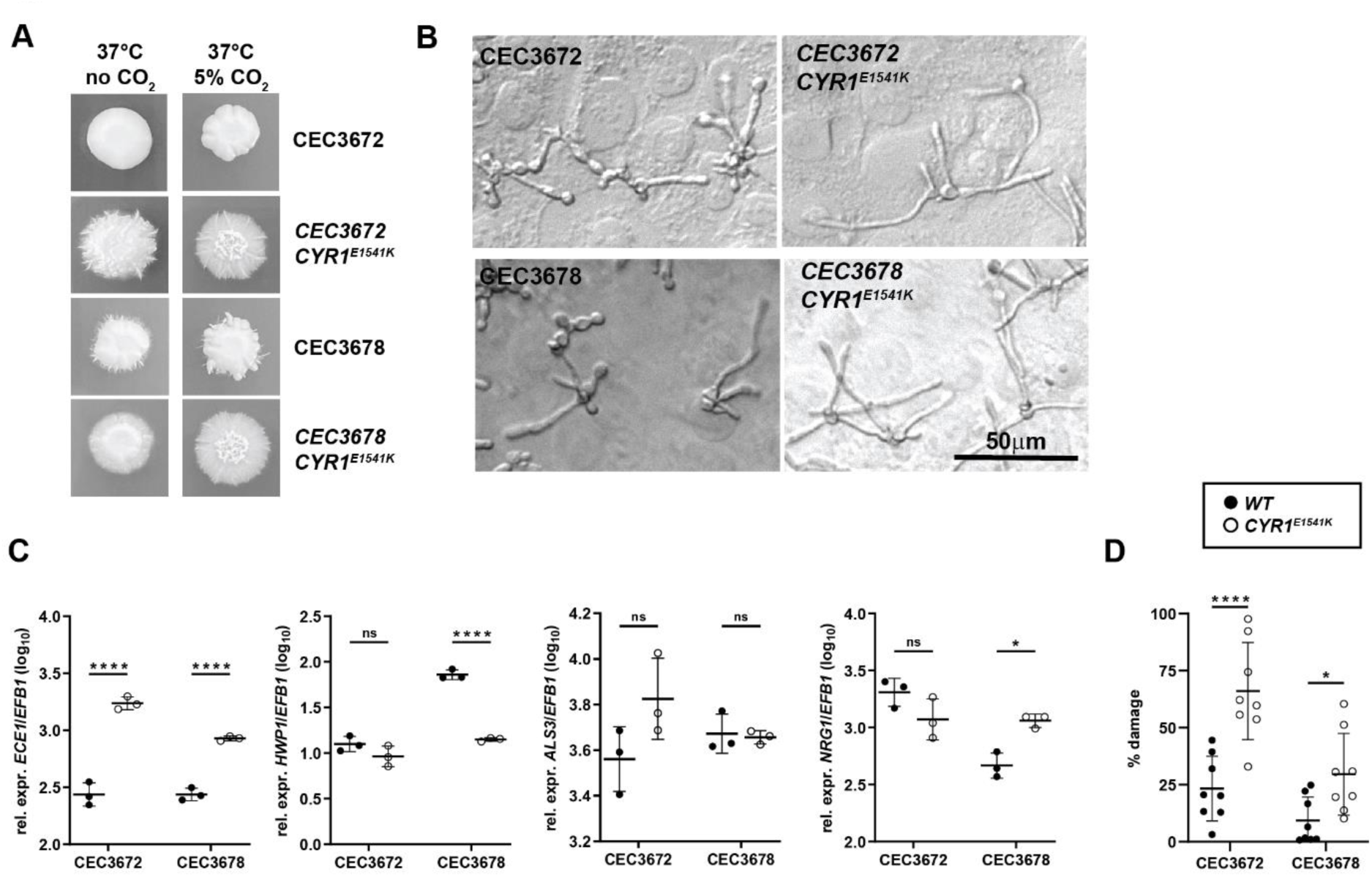
The capacity of the hyperactive *CYR1*^E1541K^ point mutation to de-repress filamentation and hyphae-associated gene expression in low-virulent *C. albicans* is strain-independent. *C. albicans* strains CEC3672, CEC3672_*CYR1*^E1541K^, CEC3678 and CEC3678_*CYR1*^E1541K^ were assessed for their phenotype *in vitro*. **A.** Colony morphology of the strains grown on Spider agar for 5 days at 37 °C**. B.** Filamentation of the strains put in contact with a monolayer of TR146 keratinocytes for 3.5 hours at 37°C, 5% CO_2_. C. *ECE1*, *HWP1*, *ALS3* and *NRG1* expression levels by the fungal strains after exposure to TR146 keratinocytes for 24 hours at 37°C, 5% CO_2_. Each symbol represents one sample. Data are from one representative out of two independent experiments with three samples per strain each. The mean ± SD is indicated. **D.** LDH release from TR146 keratinocytes after exposure to the fungal strains for 24 hours at 37°C, 5% CO_2_. Each symbol represents one well. Data are from one representative out of two independent experiments. The mean ± SD is indicated. Statistical significance was determined using unpaired Student’s *t* test comparing each mutant with the corresponding parental strain. *p<0.05, **p<0.01, ***p<0.001, ****p<0.0001. **See also Figure S6**

Together, the results from these *in vitro* and *in vivo* experiments indicate that the hyperactive variant of Cyr1 had indeed the potential to derepress some virulence traits under strong hyphae-inducing conditions such as those used for our *in vitro* assays, but it was insufficient to fully overcome the low-virulent nature of strain 101 under within-host conditions, independently of the functionality of *BRG1*.

### The capacity of the hyperactive *CYR1*^E1541K^ point mutation to de-repress filamentation and hyphae-associated gene expression in low-virulent *C. albicans* is strain-independent

To obtain further insights into the impact of the strain background on the potential of the *CYR1*^E1541K^ mutation to modulate *C. albicans* virulence, we turned to other low-virulent isolates. Strains CEC3672 and CEC3678 (38) both carry two full-length *BRG1* alleles, including the C-terminal extension, encoding Brg1 proteins almost identical to those encoded in the SC5314 genome (**Suppl. Figure S1**). These isolates exhibit a low-virulent phenotype closely comparable to that of strain 101 regarding filamentation, damage induction in keratinocytes, eliciting damage-associated cytokine release and colonization dynamics and efficiency of mice oral mucosa (13). Introducing the *CYR1*^E1541K^ point mutation in these low-virulent isolates induced filamentation as we previously observed for strain 101, both on Spider agar and when placed in contact with keratinocytes (**Figure 5A, B**, compare to **Figure 4A, B, H, I**). Likewise, the expression of hyphae-associated genes was enhanced, although not as consistently as observed in case of strain 101 (**Figure 5C**, compare to **Figure 4C, J** and **Suppl. Figure S5B, S6B**). Keratinocyte damage was also increased in response to CEC3672 and CEC3678 carrying the hypervirulent *CYR1* variant in comparison to the corresponding parental strains, whereby the degree of the effect was strain dependent (**Figure 5D**, compare to **Figure 4D, K**). Together, these data indicate that the Glu-to-Lys mutation in Cyr1 rendered the adenylate cyclase hyperactive (37), which led to de-repression of filamentation and enhanced expression of hyphae-associated genes in multiple normally low-virulent isolates, although the point mutation was insufficient to induce full-blown virulence characterized by induction of high cellular damage and elicitation of an inflammatory response at the host interface, as inferred from our results with the 101_*CYR1*^E1541K^ strain.

### The Ras1/cAMP-PKA pathway promotes virulence in intrinsically low-virulent strains

Although the spontaneously acquired Cyr1^E1541K^ mutation did not explain the entire high-virulent phenotype of strain 101_*BRG1*^FL^/*BRG1*^FL^(*), it has a remarkable capacity to alter the phenotype of several low-virulent isolates. We wondered whether the modulation of the cAMP pathway was a unique event, or may occur more frequently. Prolonged growth on Spider agar spawned several filamentation events in both strain 101 as well as the 101_*BRG1*^FL^/*BRG1*^FL^ mutant (**Figure 6A**). Clones generated from these spontaneously outgrown colonies confirmed the hyperfilamentous phenotype (**Figure 6B**). We then performed whole genome sequencing of 5 of the isolated clones to identify mutations in comparison to their parental strains. Interestingly, all identified mutations were in *RAS1* (**Figure 6C, Suppl. Table S2**), a gene encoding a GTPase that mediates Cyr1 activation in response to serum and N-acetylglucosamine stimulation through interaction with the N-terminal Ras Association domain of Cyr1 (39). Several of the identified mutations have been previously reported to render *C. abicans* and/or human Ras hyperactive (40–42). Together the data demonstrates that genes of the cAMP-PKA pathway are hotspots for spontaneously acquired mutations that confer pronounced virulence traits to *C. albicans*.

**Figure 6.**
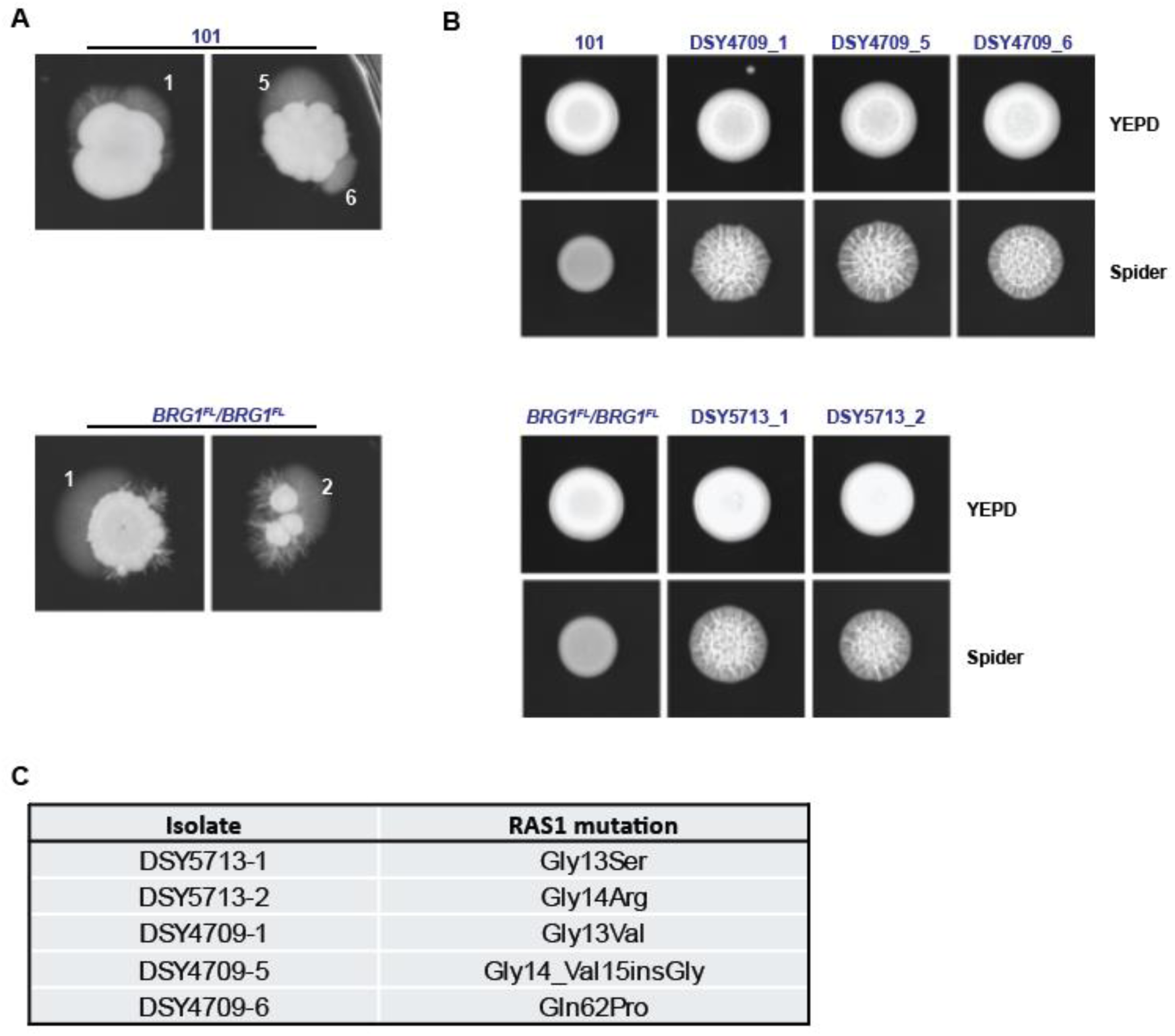
The Ras1-cAMP pathway promotes virulence in intrinsically low-virulent strains. **A.** *C. albicans* strain 101 wild type and 101_*BRG1*^FL^/*BRG1*^FL^ were plated on Spider agar for 7 days at 35 °C **B.** Colony morphology of clones that were isolated from spontaneously outgrown filaments (A) when grown on YEPD or Spider agar, as indicated. **C.** Mutations in *RAS1* identified in the clones shown in (B).

## DISCUSSION

*Candida albicans* is a prominent human pathobiont associated with diseases that vary in severity from mild superficial forms to fatal systemic infections. Despite the pathogenic potential of *C. albicans*, in a first place it is a harmless colonizer of mucosal tissues that does not damage the host, but instead rather enhance the hosts resistance to infection (5). The precise mechanisms enabling the *C. albicans* to thrive as a resident of the mycobiota remain to be fully clarified. Studying low virulence isolates and their interaction with the host has advanced the understanding of how *C. albicans* establishes colonization and avoids tissue damage and inflammation to preserve homeostasis (13, 14, 25). Here, we explored how the transcription factor Brg1 and the Cyr1 adenylate cyclase modulate virulence properties of the low-virulent*C. albicans* strain 101. While introduction of allele variants of the *BRG1* and *CYR1* genes into strain 101, either indivually or in combination, increased the strain’s virulence properties *in vitro*, it was not sufficient to render strain 101 virulent in the oral mucosal tissue of experimentally colonized mice. These findings underscore the tight restriction of the virulence gene network in *C. albicans* that guarantees homeostatic colonization in the immunocompetent host. Our data also demonstrate that filamentation and hyphae-associated gene expression under *in vitro* conditions is insufficient to predict fungal virulence *in vivo*, further highlighting the intricate regulation of fungal virulence traits at the host interface.

The newly assembled high quality genomic sequence of strain 101 enabled us conducting a comparative genome analysis of strains SC5314 and 101, which drew our attention to a rare truncated *BRG1* allele in strain 101. Brg1 regulates the *NRG1* hyphal repressor gene via mRNA destabilization (11) and chromatin remodelling (43). Our previous findings that *NRG1* expression levels are decisive for the low-virulent phenotype of strain 101 (14) led us to explore the functionally of truncated *BRG1* allele. Indeed the truncated *BRG1* allele was functionally defective. Replacing the truncated *BRG1* allele of strain 101 with a copy of the functional allele, leading to strain 101 having two full-length, functional alleles, resulted in modest increase of virulence *in vitro*, however insufficient to increase virulence in the oral mucosa *in vivo*. These data is consistent with previous reports of allelic differences in individual genes capable to modulate *C. albicans* pathogenicity (44, 45). The lack of full induction of virulence may at least in part be explained by the complex regulation of Brg1 (46, 47). We did only replace the dysfunctional *BRG1* allele with the functional allele, without directly modulating the expression levels of the gene. We did only replace the dyfunctional BRG1 ORF with the full length BRG1 ORF from strain 101, without directly modulating the expression levels. This ORF is almost identical to one of the two ORFs encoded by the first BRG1 allele in strain SC5314 but differs by several aminoacids from that encoded by the second allele. We cannot exclude that the two SC5314 Brg1 proteins have different functionalities that are required for high virulence.

Aiming to explain the acquired virulence phenotype of 101_*BRG1^FL/FL^*(*), reminiscent of the high-virulent strain SC5314, we identified a gain-of-function mutation in Cyr1, Cyr1^E1541K^, previously described to render Cyr1 hyperactive and thereby to promote constitutive filamentation of strain SC5314, even under non-hyphae-inducing conditions (37). However, in the background of strain 101, the Cyr1^E1541K^ point mutation was insufficient to fully derepress the virulent traits, irrespective of the *BRG1* status of the strain, indicating that other, so far unidentified mutations may also contribute to the conversion of a low in a high virulent strain. As a matter of fact, 101_*BRG1^FL/FL^*(*) contains in addition to the Cyr1^E1541K^ point mutation eight additional non-synonymous changes in ORFs (**Suppl. Table S1**), which may contribute to the low to high virulence of this isolate.

The *CYR1*^E1541K^dependent increase in virulence was also observed in other low-virulent strains of *C. albicans*, albeit to variable degrees. As such, the consequences of the *CYR1*^E1541K^ mutation appear to be strain-dependent. Strain-specific regulation of the activity of Cyr1 may include the availability of ATP (the substrate for the generation of cAMP), which is regulated by mitochondrial activity, or differential expression of proteins such as Pde2 (17).

We also observed a difference in the impact of *CYR1*^E1541K^ on fungal virulence *in vitro* and *in vivo*, indicating that the environmental cues inducing filamentation and the hyphae-associated virulence program differ in cell culture and inside mammalian tissues. As such, the degree to which the cAMP-PKA pathway is required for filamentation may also differ under *in vivo* and *in vitro* conditions (48). In the mucosal tissue, the treshold to de-repress the low-virulence program is particularly high and requires an intricate sequence of events that unfold over extended periods of time (Fróis-Martins et al., under revision). Cyr1 hyperactivation appears not to be strong enough to override the negative regulation of filamenation and hyphae-associated genes by Tup1 and Nrg1 (16). *C. albicans* is able to colonize different types of host tissue such as the gut that present distinct environmental context (1, 49). For instance, bacterial peptidoglycan-like molecules, which are abundant in the intestine, were shown to induce filamentation by directly binding to Cyr1 (50). We only evaluated the oral mucosa, therefore we cannot exclude a different effect of the Cyr1 point mutation on the fungal phenotype in other tissue comparments.

In addition to the initially identified spontaneous high-virulent mutant 101_*BRG1^FL/FL^*(*), we recovered additional spontaneous mutants outgrowing from strain 101, which all exhibited mutations in the N-terminal part of the *RAS1* gene known to freeze the protein in a permanently active state (51, 52) Ras1 acts upstream of Cyr1 to induce filamentation (17). In addition, Ras1 can also induce filamentation independently of Cyr1 by activating Cdc24, which in turn activates the Cph1 transcription factor involved in the filamentation program (53). The observation that all spontaneously developed variants exhibited mutations affecting the cAMP-PKA pathway raises important questions regarding the driving force inducing these mutations. An environmental cue specific to the conditions under which the fungus was cultured may specifically target the cAMP-PKA pathway, or the cAMP-PKA signaling cascade may serve as a preferential target due to its central role in facilitating environmental adaptation without significantly jeopardizing the fungus’ survival.

Together, we demonstrate that strain 101 tightly regulates fungal virulence traits. Additional allele variants may contribute to the inherently low-virulent properties, and replacing an individual gene turned out to be insufficient to overcome the firmly anchored strain-intrinsic colonizer program, which supports the maintenance of homeostasis in the colonized mucosal tissue. In addition to the strain-intrinsic properties of a strain, the functional state of a strain also depends on the environmental conditions. When exposed to hyphae-inducing conditions, such as those characteristic of the macrophage phagosome or in the epithelium of the IL-17 deficient host, even low-virulent strains can gradually express increased virulence features (54–56) (Fróis-Martins et al., under revision). We observed the similar phenomenon when growing strain 101 on Spider agar thereby further demonstrating the strong adaptatability of *C. albicans* to its environment.

## MATERIALS AND METHODS

### Fungal strains

Strains used in this study are listed in **Table 1**. All strains were maintained on YPD agar for short term and in glycerol-supplemented medium at -80°C for long-term storage. Cultures were inoculated at OD_600_ = 0.1 in YPD medium (using a pre-culture to adjust the OD_600_) and grown at 30°C and 180 rpm for 15-18 hours. At the end of the culture period, yeast cells were washed in PBS and their concentration was determined by spectrophotometry, whereby 1 OD_600_ = 10^7^ yeast cells.

**Table 1.** C. albicans strains used in this study.

| Strain | Collection # | genotype | Parental | reference |
| --- | --- | --- | --- | --- |
| SC5314 | CA1 | Clinical isolate | n/a | (57) |
| 101 | DSY4709<br>CA117 | Clinical isolate | n/a | (58) |
| CEC3672 | CA327 | Clinical isolate | n/a | (38) |
| CEC3678 | CA328 | Clinical isolate | n/a | (38) |
| SC5314_*brg1*Δ/Δ | DSY5875<br>CA329 | *arg4*Δ/*arg4*Δ, *leu2*Δ/*leu2*Δ, *his1*Δ/*his1*Δ, *URA3*/iΔ, *IRO1*/*iro1*Δ, *brg1*Δ::*HIS1*, *brg1*Δ::*LEU2* | SC5314 | this study |
| SC5314_*BRG1*/BRG1 | DSY5876<br>CA330 | *arg4*Δ/*arg4*Δ, *LEU2*/*leu2*Δ, *HIS1*/*his1*Δ, *URA3*/*ura3*Δ, *IRO1*/*iro1*Δ | DSY5875 | this study |
| SC5314_*brg1*Δ/*BRG1*<sup>FL</sup> | DSY5877<br>CA331 | *arg4*Δ/*arg4*Δ, *leu2*Δ/*leu2*Δ, *his1*Δ/*his1*Δ, *URA3*/*ura3*Δ, *IRO1*/*iro1*Δ, *brg1*Δ::*BRG1*<sup>FL</sup>::*SAT1*, *brg1*Δ::*LEU2* (FL allele of strain 101) | DSY5875 | this study |
| SC5314_*brg1*Δ/*BRG1*<sup>TRUNC</sup> | DSY5880<br>CA334 | *arg4*Δ/*arg4*Δ, *leu2*Δ/*leu2*Δ, *his1*Δ/*his1*Δ, *URA3*/*ura3*Δ, *IRO1*/*iro1*Δ, *brg1*Δ::*BRG1*<sup>TRUNC</sup>::*SAT1*, *brg1*Δ::*LEU2* (TRUNC allele of strain 101) | DSY5875 | this study |
| SC5314_*BRG1*/BRG1<sup>TRUNC</sup> | DSY5720<br>CA273 | *BRG1*/ *BRG1*<sup>TRUNC</sup> ::*NAT1* | SC5314 | this study |
| 101_*BRG1*<sup>FL</sup>/BRG1<sup>TRUNC</sup> | DSY5710<br>CA267 | *BRG1*<sup>FL</sup>/BRG1<sup>TRUNC</sup> ::*NAT1* | 101 | this study |
| 101_*BRG1*<sup>FL</sup>/BRG1<sup>FL</sup> | DSY5713<br>CA270 | *BRG1*<sup>FL</sup>/BRG1<sup>FL</sup> ::*NAT1* | 101 | this study |
| 101_*BRG1*<sup>FL</sup>/BRG1<sup>FL</sup>(*) | DSY5722<br>CA271 | *BRG1*<sup>FL</sup>/BRG1<sup>FL</sup> ::*NAT1*, *CYR1*/*CYR1*<sup>E1541K</sup> | DSY5713 | this study |
| 101_*CYR1*<sup>E1541K</sup> | DSY5731<br>CA279 | *CYR1*/*CYR1*<sup>E1541K</sup>, *ENO1*::pV1025 | 101 | this study |
| 101_*CYR1*<sup>E1541K</sup>/*CYR1*<sup>E1541K</sup> (#2) | DSY5730<br>CA295 | *CYR1*<sup>E1541K</sup>/*CYR1*<sup>E1541K</sup>, *ENO1*::pV1025 | DSY5731 | this study |
| 101_*CYR1*<sup>E1541K</sup>/*CYR1*<sup>E1541K</sup> (#2) | DSY5732<br>CA296 | *CYR1*<sup>E1541K</sup>/*CYR1*<sup>E1541K</sup>, *ENO1*::pV1025 | DSY5731 | this study |
| 101_ *BRG1*<sup>FL</sup>/*BRG1*<sup>FL</sup><br>*CYR1*<sup>E1541K</sup> | DSY5862 | *BRG1*<sup>FL</sup>/*BRG1*<sup>FL</sup>::*NAT1*,<br>*CYR1*<sup>E1541K</sup>, *ENO1*::pDS2168 | DSY5713 | this study |
| 101_ *BRG1*<sup>FL</sup>/*BRG1*<sup>FL</sup><br>*CYR1*<sup>E1541K</sup> | DSY5863 | *BRG1*<sup>FL</sup>/*BRG1*<sup>FL</sup>::*NAT1*, *CYR1*<sup>E1541K</sup>,<br>*ENO1*::pDS2168 | DSY5713 | this study |
| CEC3672_ *CYR1*<sup>E1541K</sup> | DSY5791<br>CA299 | *CYR1*/*CYR1*<sup>E1541K</sup>, *ENO1*::pV1025 | CEC3672 | this study |
| CEC3678_ *CYR1*<sup>E1541K</sup> | DSY5774<br>CAC303 | *CYR1*/*CYR1*<sup>E1541K</sup>, *ENO1*::pV1025 | CEC3678 | this study |
| DSY4709_1 (*) | DSY5840 | *RAS1*/*RAS1*<sup>G14R</sup> | 101 | this study |
| DSY4709_5 (*) | DSY5842 | *RAS1*/*RAS1*<sup>G13S</sup> | 101 | this study |
| DSY4709_6 (*) | DSY5844 | *RAS1*/*RAS1*<sup>G14V</sup> | 101 | this study |
| DSY5713_1 (*) | DSY5846 | *BRG1*<sup>FL</sup>/*BRG1*<sup>FL</sup>::*NAT1*,<br>*RAS1*/*RAS1*<sup>G14V15insG</sup> | DSY5713 | this study |
| DSY5713_2 (*) | DSY5848 | *BRG1*<sup>FL</sup>/*BRG1*<sup>FL</sup>::*NAT1*,<br>*RAS1*/*RAS1*<sup>Q62P</sup> | DSY5713 | this study |

**Table 2:**
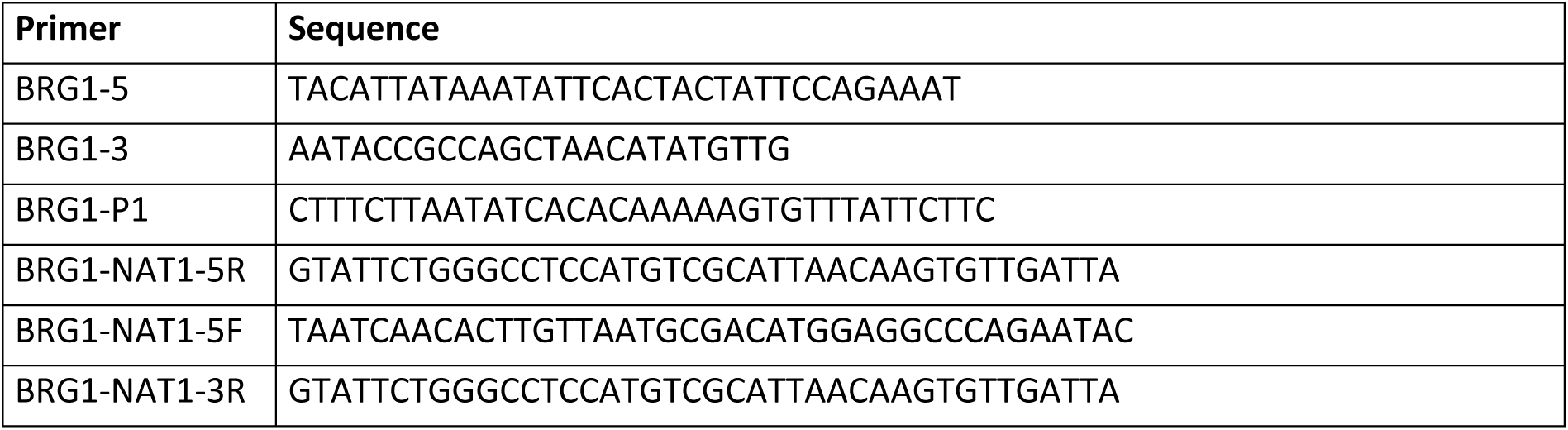
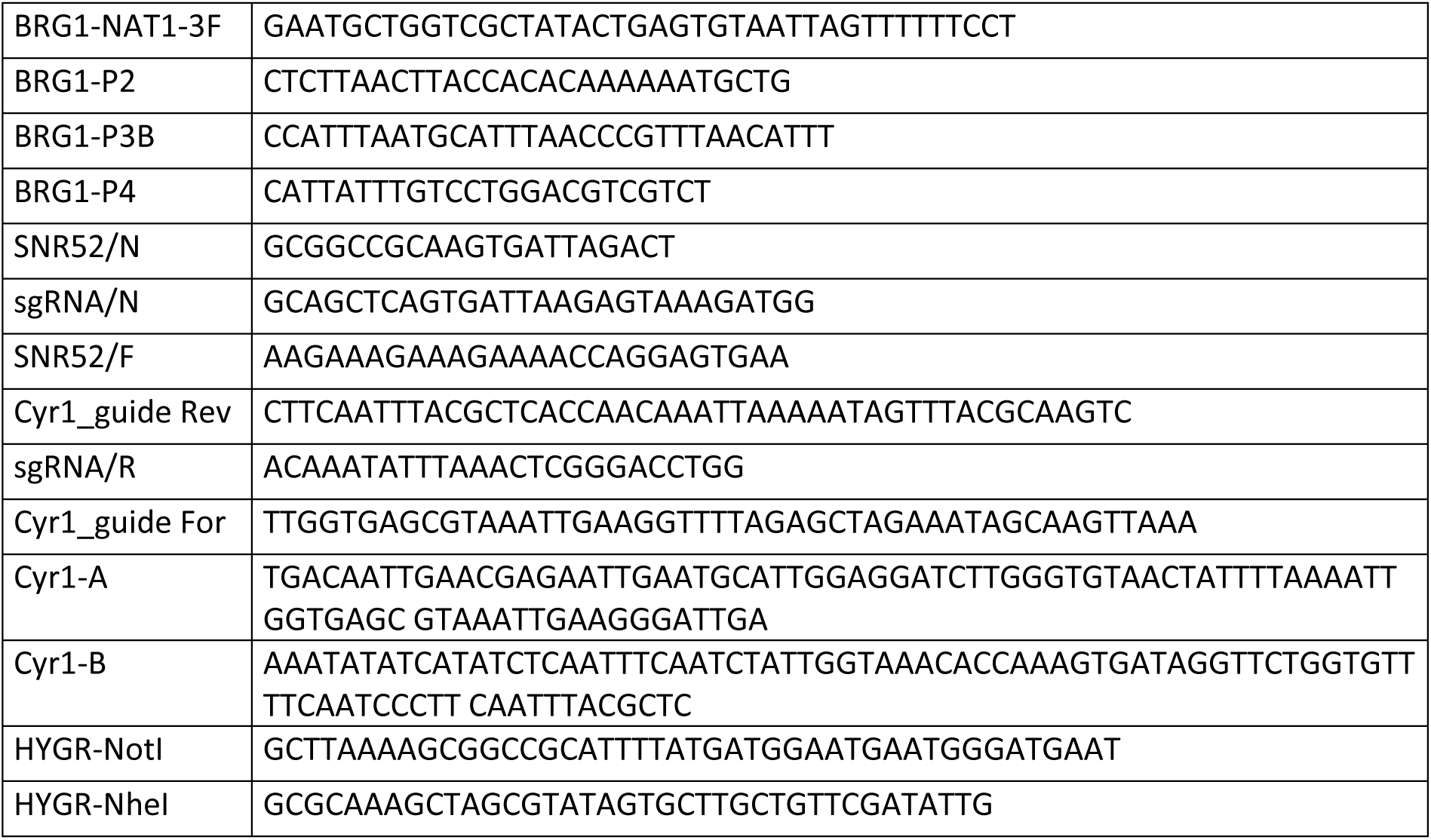
primer list.

### Identification of the truncated *BRG1* allele

PCR of *BRG1* alleles on strain 101 was accomplished with primers BRG1-5 and BRG1-3 and analysed by gel electrophoresis on 1% agarose. The *BRG1* truncated allele (*BRG1*^TRUNC^) shows a reduced size (1.3 kb) as compared to the full-length *BRG1* allele (1.5 kb; *BRG1*^FL^).

### Mutant generation

*BRG1*^TRUNC^ and *BRG1*^FL^ revertants in the SC5314 background:

*BRG1* revertants were constructed with a *brg1*Δ/Δ mutant available from the Hohmann mutant collection (DSY5875) (59). A PCR fusion strategy was used to amplify *BRG1* alleles fused to the *NAT1* selection marker. The *BRG1* alleles with 500 bp promoter from 101 were first amplified with primers BRG1-P1 and BRG1-NAT1-5R (overlapping primer with *NAT1*). Next, the NAT1 marker was amplified from pJK795 with BRG1-NAT1-5F and BRG1-NAT1-3R. Finally, the *BRG1* dowstream sequence (500 bp) was amplified with primers BRG1-P2 and BRG1-NAT1-3F (overlapping primer with *NAT1*). After PCR fragment purification, equimolar amounts of the 3 fragments were used with nested primers BRG1-P3B and BRG1-P4 in a fusion PCR to results in a 4 kb fragment. This fragment was used to transform DSY5875 in order to replace the deleted *BGR1* loci with 101 *BRG1* alleles. This was achieved by a RNA-protein complexes (RNPs) approach as reported in Grahl *et al.* (60) that employs reconstituted purified Cas9 protein in complex with scaffold and gene-specific guide RNAs. A gRNA specific for the *BRG1* promoter (TGGGTGTAGAGAAACGATGT) upstream of the fusion PCR construct was designed in silico using Geneious Prime and obtained from IDT (Integrated DNA Technologies, Inc.) as CRISPR guide RNA (crRNA), which contains 20-bp homologous to the target gene fused to the scaffold sequence. RNPs were created using the Alt-R CRISPR-Cas9 system from IDT. Briefly, The *BRG1* crRNA and tracrRNA (a universal transactivating CRISPR RNA) were first dissolved in RNase-free distilled water (dH_2_O) at 100 µM and stored at -80°C. The complete guide RNA was generated by mixing equimolar concentrations (4 µM final) of the gene-specific crRNA and tracrRNA to obtain a final volume of 3.6 µl per transformation. The mix was incubated at 95°C for 5 min and cool down to room temperature. The Cas9 nuclease 3NLS (60 µM stock from IDT) was diluted to 4 µM in dH_2_O at a volume of 3 µl per transformation. RNPs were assembled by mixing guide RNAs (3.6 µl of gene-specific crRNA/tracrRNA) with 3 µl of diluted Cas9 protein, followed by incubation at room temperature for 5 min. Transformation of *C. albicans* was carried out by electroporation and used 6.6 µl of gene-specific RNPs, 40 µl of *C. albicans* cells and 1-2 µg of the *BRG1* fusion PCR (up to 3.4 µl volume). Transformants were selected onto YEPD plates with nourseothricin (200 µg/mL) for 3 days at 35°C. After transformants selection, identification of the specific 101 *BRG1* alleles introduced in DSY5875 were verified by targeted sequencing analysis. DSY5877 contained the *BRG1*^FL^ allele replacing the *HIS3* selective marker in DSY5875, while DSY5880 contained the *BRG1*^TRUNC^ allele replacing the *LEU2* selective marker in the same strain.

A wild type *BRG1* allele was also replaced by *BRG1*^TRUNC^ allele in SC5413. For this, construction of the repair fragment followed the above-mentioned fusion PCR approach. Two guides specific for *BRG1* in SC5314 including 5-BRG1-guide-SC (TGGGAAAGAAGAGATTTACA) and 3-BRG1-guide-SC (TTTAAAAAGCATCAATCAAT) were used in the RNP complexes construction. A minor adaptation was that the RNPs were concentrated by 2-fold and mixed each (3,3 µl) with the *BRG1* repair fragment for electroporation. Transformants were analysed by targeted sequence analysis and yielded DSY5720.

*BRG1*^TRUNC^ and *BRG1*^FL^ revertants in the 101 background:

In order to replace specific *BRG1* alleles in the 101 background, two different guides fusion were designed including 5-BRG1-guide-mut-allele (TATTTATAATGTAACAAGTA) and 3-BRG1-guide-mut-allele (CTGCACTGAGATATAAAAAG) that were used in the RNP complexes construction. The RNPs were concentrated by 2-fold and mixed each (3,3 µl) with the *BRG1* repair fragment for electroporation. Transformants were analysed by targeted sequence analysis and yielded DSY5710 and DSY5713 containing each *BRG1*^FL^/*BRG1*^TRUNC^ alleles and *BRG1*^FL^/*BRG1*^FL^ alleles, respectively.

Generation of Cyr1 ^E1541K^ mutants:

The introduction of the E1541K mutation in Cyr1 was realized with a transient guide RNA expression following published protocols (61) . First, the guide expression was constructed with 2 overlapping PCRs obtained with primers pairs SNR52/F-Cyr1_guide Rev and sgRNA/R-Cyr1_guide For using pV1093 (62) as template. Fusion PCR between the two fragments was performed with primers SNR52/N and sgRNA/N. Next, the repair fragment (containing the Cyr1 ^E1541K^ mutation without PAM sites) was obtained by mixing primers Cyr1-A and Cyr1-B followed by self priming and 20 PCR cycles. The two PCR were then mixed with pV1025 previously digested by *Sac*I and *Kpn*I to transform strains 101, CEC3672 and CEC3678 (14) by electroporation. Transformants were selected onto YEPD plates with nourseothricin (200 µg/mL) for 3 days at 35°C. Targeted sequencing of transformants yielded heterozygous (DSY5731, DSY5791, DSY5774) and homozygous (DSY5730, DSY5732) Cyr1 ^E1541K^ mutants.

In order to introduce the E1541K mutation in the background of isolate DSY5713, pV1025 was first modified by replacing *SAT1* by *HYGR* (hygromycin resistance). This was achieved by PCR amplification of *HYGR* from PYM70 (63) with primers HYGR-NotI et HYGR-NheI and insertion in pV1025 at the *Not*I and *Nhe*I restriction site thus yielding pDS2168. This plasmid was digested by *Kpn*I and *Eco*RV in the above-described transformation procedure. After transformants selection, targeted sequencing yielded heterozygous Cyr1 ^E1541K^ mutants DSY5862 and DSY5863.

### Genome assembly and annotation of strain 101

High quality genomic DNA was prepared according to published procedures (64) . High molecular weight DNA was sheared with Megaruptor (Diagenode, Denville, NJ, USA) to obtain 10-15 kb fragments. After shearing the DNA size distribution was checked on a Fragment Analyzer (Advanced Analytical Technologies, Ames, IA, USA). The sheared DNA (1.8 µg) was used to prepare a SMRTbell library with the PacBio SMRTbell Express Template Prep Kit 2.0 (Pacific Biosciences, Menlo Park, CA, USA) according to the manufacturer’s recommendations. The resulting library was size-selected on a Blue Pippin system (Sage Science, Inc. Beverly, MA, USA) for molecules larger than 6 kb. It was sequenced with v2.2/v2.0 chemistry and adaptive loading on a PacBio Sequel II instrument (Pacific Biosciences, Menlo Park, CA, USA) with 30h movie length, pre-extension time of 2h using one SMRT cell 8M. Genome assembly was conducted with hifiasm (v0.16.1-r375) (65), and the resulting contigs were benchmarked against the *C. albicans* SC5314 reference (A22-s07-m01-r145) using MUMmer (v3.0) (66), QUAST (67) and *BUSCO* (v5.4.2) (68) to assess completeness, contiguity and gene content. Ab initio gene prediction was performed with AUGUSTUS (v3.4.0) (69), and annotations were refined by mapping to closest orthologs from the OMA Knowledgebase using OMA Fast Mapping (70). Protein-coding gene models underwent manual curation in Geneious Prime (www.geneious.com) to resolve ambiguities and validate exon–intron boundaries. Non-coding RNA loci were identified with Infernal (v1.1.4) (71) against the Rfam database (v14.8) (68). Genome feature files (GFF) and sequence files (FASTA) were manipulated and formatted using EMBOSS (v6.6.0) (72) and integrated into the final annotation with MAKER (v2.31.9) (73). The complete diploid assembly and raw sequencing reads are available under NCBI BioProject PRJNA923600.

### Genome-wide mutant analysis

Spontaneous filamenting mutants were subjected to genome sequence analysis by comparison with the newly genome-annotated genome of strain 101. Genomic DNA from isolates DSY5722, DSY5846 and DSY5848 (DSY5713 as parent) and isolates DSY5840, DSY5842 and DSY5844 (101 as parent) was first obtained by a spheroplasting method (74). Genomic libraries were prepared from 100 ng of DNA with the Illumina DNA Prep Kit (Illumina) and Unique Dual Indexed oligonucleotides (IDT). PCR amplification was performed for 5 cycles. Libraries were quantified by a fluorimetric method and their quality assessed on a Fragment Analyzer (Agilent Technologies). Sequencing was performed as a 2x150 cycles paired run either on a MiSeq v2 flow cell (DSY5713 and DSY5722) or a NovaSeq 6000 v1.5 flow cell (Illumina) (DSY5846, DSY5848, DSY5840, DSY5842 and DSY5844). Sequencing data were demultiplexed using the bcl2fastq2 Conversion Software (v. 2.20, Illumina). Data can be obtained under Bioproject PRJNA1147312.

Single nucleotide polymorphism (SNP) analysis was conducted to compare the isolates DSY5713 and spontaneous filamenting mutant DSY5722 using the SC5314 reference genome (A22-s07-m01-r145) as a reference. Raw sequencing reads from both isolates were aligned to the SC5314 genome using Bowtie2 (v2.3.4.1) (75) . Variant calling was performed with FreeBayes (v1.2.0) (76), a haplotype-based variant detector, to identify SNPs and small insertions or deletions. The resulting variant call format (VCF) files from both isolates were normalized and filtered using SAMtools (v1.10) (77) and VCFtools (v0.1.15) (78) to enable accurate comparative analysis. Finally, the functional impact of the identified variants was assessed using SnpEff (v4.3) (79), which annotated the variants based on gene models and predicted their potential effects on protein function. SNPs specific to DSY5722 were filtered from those present in the parent DSY5713 (**Suppl. Table S1**).

SNP analysis from isolates DSY5846, DSY5848, DSY5840, DSY5842 and DSY5844 was performed with CLC Genomics Workbench (v24) and the variant detection tools using the annotated genome from isolate 101. After read mapping to the 101 genome, variants differing from those present in DSY5713 and DSY5722 were filtered using the variant filtering tool in CLC Genomics Workbench (**Suppl. Table S2**).

#### Growth curve assays

*C. albicans* was inoculated at OD_600_ = 0.1 in 200 μl YPD or F12 cell culture medium (see below) in a 96well plate and grown in Synergy H1 plate reader (Bio Tek) for 24 hours at 30°C with double orbital shaking (3 mm amplitude) and OD600 measurements every 15 minutes that were preceded by a 10 s linear shaking pulse (1 mm amplitude).

#### Colony morphology assays

For assessing filamentation on spider agar (10 g D-Mannitol, Sigma, Cat.no. M4125, 10g nutrient broth no.1, Sigma, Cat.no. 70122, 2g K_2_HPO_4,_ Sigma, Cat.no. 795496 and 13.5g agar, Sigma, Cat.no. A1296 per 1L H_2_O), 3 µl of a yeast cell suspension at 10^5^ cells/ml were spotted and plates were incubated for specific timing and tempratures as indicated in Figure legends. In some experiments, plates were incubated in presence of 5% CO_2_. Filamentation was assessed on YEPD agar (yeast Extract Peptone Dextrose (YEPD) (1% Bacto peptone, Difco Laboratories, Basel, Switzerland), 0.5% yeast extract (Difco) and 2% glucose (Fluka, Buchs, Switzerland) 2% agar (Difco) and YCB/BSA agar (1.17% Yeast Carbon Base (Becton Dickinson), 2% BSA (Sigma). Overnight cultures in YEPB were diluted in water to obtain single colonies. Plates were incubated at 35°C for 5- to 7 days. YCB/GlcNAC contained 1.17% Yeast Carbon Base and 2% N-Acetyl-D-glucosamine (Sigma) and 2% agar when indicated.

#### Keratinocyte cell culture

The human oral keratinocyte cell line TR146 (80) was grown in DMEM medium (Sigma, Cat.no. D5796) supplemented with 10% FCS, 1% Penicillin and 1% Streptomycin at 37°C and 5% CO2. For RNA isolation and filamentation assay, cells were seeded at 1 x 10^5^ cells/well in 24-well tissue culture plates and for cell damage assay at 4 x 10^4^ cells/well in 96-well tissue culture plates, respectively, and grown to confluent monolayers for 2 days prior to infection as described below for the individual assays. 1 day prior to the experiment, the DMEM medium was replaced by F12 medium (Hams’s Nutrient Mixture F12 medium (Gibco, Cat.no. 21765029) supplemented with 1% FCS).

#### Filamentation assay

Monolayers of TR146 cells in 24-well tissue culture plates prepared as described above were infected with 5 x 10^4^ yeast cells per well in a 24-well plate and incubated for 3.5 hours at 37°C and 5% CO_2_. Cells were fixed in 2 % PFA a for 20 minutes at 4°C. The PFA was then exchanged by PBS for imaging with an EVOS FL Auto microscope (Life Technologies). Filament length was determined with Image J. 20-30 filaments were analyzed per imaged fields, whereby 10 fields were imaged per well and 2 wells were analyzed per strain.

#### Cell damage assay

Damage induction in TR146 cells by *C. albicans* was performed as described (81). Briefly, cell monolayers in 96-well tissue culture plates prepared as described above were infected with 2 x 10^4^ yeast cells per well and incubated for 24 hours at 37°C and 5% CO2. Control wells were incubated with medium only or with 1% Triton-X-100 to determine 100% damage. LDH release into the supernatant was quantified with the LDH cytotoxicity kit (Roche) according to the manufacturer’s instructions.

#### RNA isolation from infected keratinocyte cultures

Monolayers of TR146 cells in 24-well tissue culture plates prepared as described above were infected with 1 x10^5^ yeast cells per well. After 24 h of incubation at 37°C and 5% CO2, the medium was removed, cell were frozen on liquid nitrogen and stored at -80°C until further processing. RNA was isolated with the RNeasy Mini Kit (Quiagen). Briefly, the cells were lyzed in RLT lysis buffer and homogenized with 0.5 mm glass beads (Sigma, Cat.no. G1277) using a Tissue Lyzer (Qiagen) 7 times for 2 minutes at 30 Hz interrupted by 30sec cooling periods on ice. Cell debris were removed by centrifugation. One volume of 75% ethanol was admixed, and each sample was transferred to a RNeasy spin column. The loaded columns were washed using RW1 buffer and RPE buffer. RNA was eluted in 30 μl RNAse-free water.

#### Animals

WT C57BL/6j mice were purchased by Janvier Elevage and kept in specific pathogen-free conditions at the Institute of Laboratory Animals Science (LASC, University of Zurich, Zurich, Switzerland). Female mice were used at 8-14 weeks inage-matched groups. Infected and uninfected animals were kept separately to avoid cross-contamination.

#### Ethics statement

All mouse experiments in this study were conducted in strict accordance with the guidelines of the Swiss Animals Protection Law and were performed under the protocols approved by the Veterinary office of the Canton Zurich, Switzerland (license number 167/2018 and 141/2021).

#### Oral colonization of mice with *C. albicans*

Mice were infected sublingually with 2.5 x 10^6^ *C. albicans* yeast cells as described (82), without immunosuppression. In brief, mice were anesthetized by injection of 100 mg/kg Ketamin and 20 mg/kg Xylazin in sterile saline i.p. administered in three doses. *C. albicans* was administered by depositing a 2.5 mg cotton ball that was soaked in 100 µl suspension at 5 x 10^7^ yeast cells/ml under the tongue for 80-90 minutes. Mice were kept on a heating mat at 35 - 37°C during the entire period of anesthesia, administered with 10 ml/kg sterile saline to stabilize the circulation, and vitamin A ointment was applied to avoid drying out of the eyes.

#### Determination of fungal burden

For determination of the fungal burden, the tongue of euthanized animals was removed, homogenized in sterile 0.05 % NP40 in H_2_O for 3 minutes at 25 Hz using a Tissue Lyzer (Qiagen) and serial dilutions were plated on YPD agar containing 100 µg/ml Ampicillin.

#### Histology

Tongue tissue was fixed in 4% PBS-buffered paraformaldehyde overnight and embedded in paraffin. Sagittal sections (9µm) were stained with Periodic-acidic Schiff (PAS) reagent and counterstained with Haematoxilin and mounted with Pertex (Biosystem) according to standard protocols. Images were acquired with a digital slide scanner (NanoZoomer 2.0-HT, Hamamatsu) and analyzed with NDP.view2.

#### RNA isolation from colonized tongue tissue

Isolation of total RNA (including fungal RNA) from murine tongue was performed using TRI reagent (Sigma-Aldrich) according to the manufacturer’s protocol. Prior to the addition of chloroform, the tongue tissue was homogenized in presence of a steel ball for 3 minutes at 25Hz followed by homogenization with 0.5 mm glass beads (Sigma) 2 times for 2 minutes at 30 Hz with a 30 second cooling period in between. Both homogenizations were performed using a Tissue Lyzer (Qiagen).

#### RT-qPCR

cDNA was generated by RevertAid reverse transcriptase (ThermoFisher Scientific, Cat.no. EP0452). Quantitative PCR was performed using SYBR green (Roche, Cat.no. 4913914001) and a QuantStudio 7 Flex instrument (Life Technologies). The primers used in this study are listed in **Table 3**. All qPCR reactions were performed in duplicates, and the relative expression (rel. expr.) of each gene was determined after normalization to *EFB1* or *ACT1* housekeeping gene transcript levels.

**Table 3.**
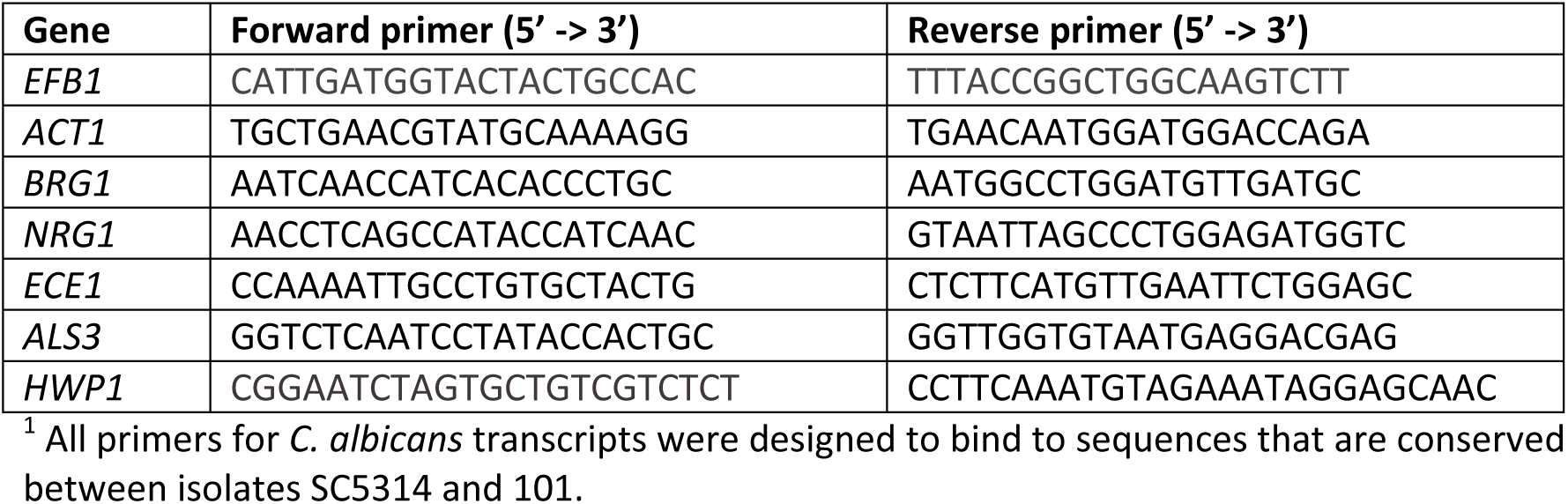
Primers used in this study for detecting C. albicans transcripts^1^.

## ACKNOWLEDGMENTS

The authors would like to thank the staff of the Laboratory Animal Service Center of University of Zürich for animal husbandry; staff of the Laboratory for Animal Model Pathology of University of Zürich for histology; the Zentrum für klinische Studien of the Vetsuisse Faculty, University of Zurich for access to equipment; Kontxi Martinez de San Vicente for foundational experiments that led to the initiation of this project; members of the LeibundGut-lab for helpful advice and discussions. This work was supported by the Swiss National Science Foundation (grant # CRSII5_173863 to S.L.L., D.S. and CdE). Work in the laboratory of CdE is supported by the Agence Nationale de Recherche (ANR-10-LABX-62-IBEID).

## AUTHOR CONTRIBUTIONS

R.F.M., D.S. and S.L.-L. designed the study and wrote the manuscript. R.F.M. and S.M. performed the experiments and analyzed the data. D.S. generated *C. albicans* mutants. V.D.T. did the bioinformatic analyses of strain 101. C.M. and C.d’E did additional bioinformatic analyses. S.L.-L., D.S. and C.d’E. oversaw the study design and data analysis, acquired funding. All authors discussed the results and commented on the manuscript.

---

**Figure S1 (related to Figure 1).**
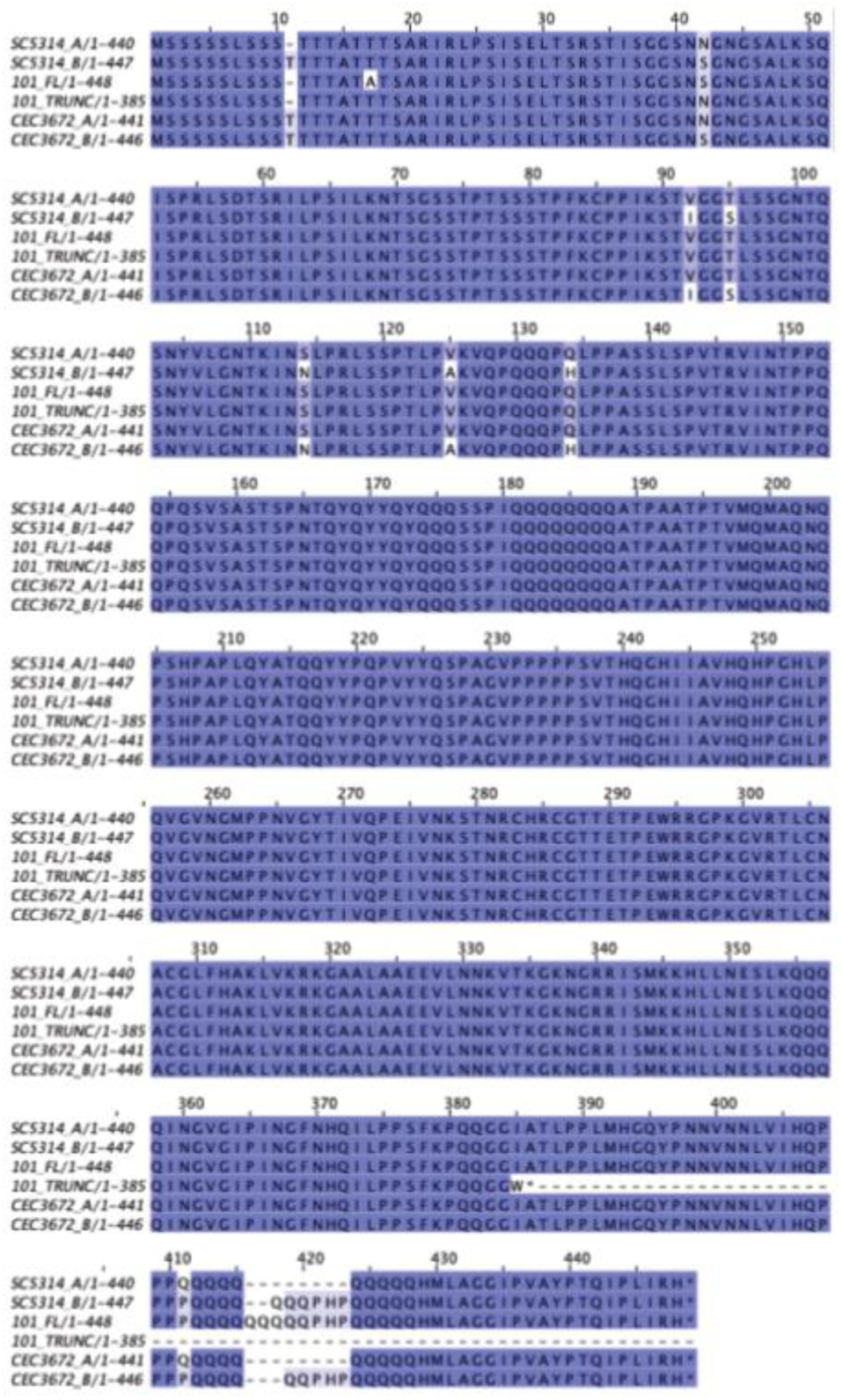
Alignment of Brg1 proteins from *C. albicans* strains SC5314, 101 and CEC3672. For strains SC5314 and CEC3672, *BRG1* haplotypes were inferred from SNP data obtained for 182 genome-sequenced strains (24) using PHASE 2.0 as previously described (Wickramasinghe *et al*., 2024). Haplotypes were subsequently edited to consider insertion and deletion events identified using GATK (24). For strain 101, *BRG1* haplotypes were obtained based on the genome assembly produced in this study. The deduced aminoacid sequences for the 6 *BRG1* haplotypes were aligned using MUSCLE (83) and colored using JalView (84). The level of conservation of the aminoacids across the 6 sequences is shown with a descending coloring ranging from dark blue to white. The aminoacid C-terminal extension shared by all Brg1 proteins and lacking in the Brg1 protein deduced from Assembly 22 of the *C. albicans* SC5314 genome is boxed in red. Data obtained for strain CEC3678 are not shown as they are identical to those obtained for strain CEC3672.

**Figure S2 (related to Figure 1).**
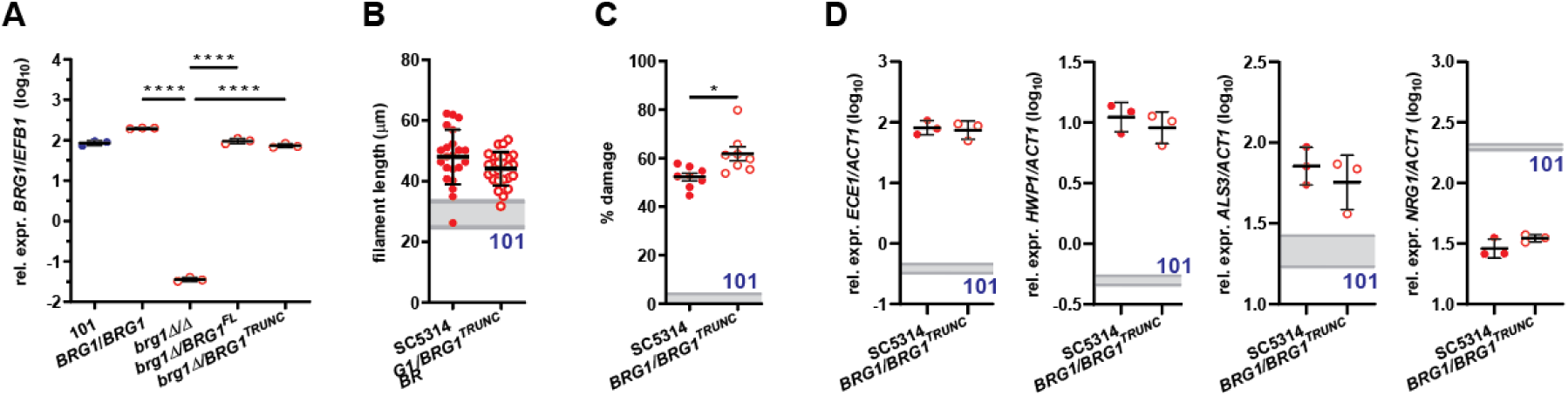
The truncated *BRG1* allele identified in the low-virulent strain 101 is unable to drive virulence in strain SC5314. A. *BRG1* expression levels in *C. albicans* strains 101 wild type, SC5314_*BRG1*/*BRG1*, SC5314_*brg1*Δ/Δ, SC5314_*brg1*Δ/*BRG1*^FL^, SC5314_ *brg1*Δ/*BRG1*^TRUNC^ after exposure to TR146 keratinocytes for 24 hours at 37°C, 5% CO_2_. Each symbol represents one sample. Data are from one representative out of two independent experiments with three samples per strain each. The mean ± SD is indicated. C.-E. *C. albicans* strains SC5314 wild type and SC5314_*BRG1*/*BRG1*^TRUNC^ were assessed for their phenotype *in vitro*. **C.** Quantification of filament length of strain put in contact with TR146 cells. Each symbol represents the mean filament length of 20 – 30 filaments per visual field. Data are pooled from two independent experiments with 10-12 visual fields analyzed per strain each. The mean ± SD is indicated. **D.** LDH release from TR146 keratinocytes after exposure to the fungal strains for 24 hours at 37°C, 5% CO_2_. Each symbol represents one well. Data are pooled from two independent experiments with 4 wells per strain each. The mean ± SEM is indicated. E. *ECE1*, *HWP1*, *ALS3* and *NRG1* expression levels by the fungal strains after exposure to TR146 keratinocytes for 24 hours at 37°C, 5% CO_2_. Each symbol represents one sample. Data are from one representative out of two independent experiments with three samples per strain each. The mean is indicated. Statistical significance was determined using one-way ANOVA (C) or student’s *t*-test (D-F).. ***p<0.001, ****p<0.0001.

**Figure S3 (related to Figure 2).**
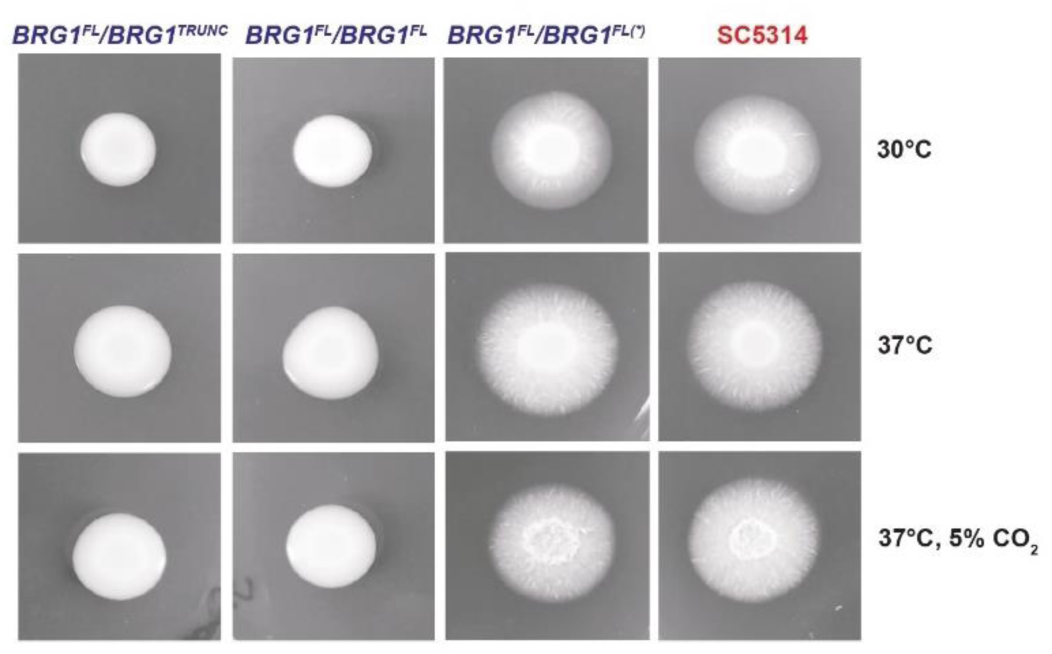
Restoring the full-length *BRG1* allele in low-virulent strain 101 is insufficient to increase the strain’s virulence. *C. albicans* strains 101_*BRG1*^FL^/*BRG1*^TRUNC^, 101_*BRG1*^FL^/*BRG1*^FL^, 101_*BRG1*^FL^/*BRG1*^FL^(*), and SC5314 wild type were assessed for their phenotype *in vitro*. **A.** Colony morphology of the strains grown on Spider agar (top and middle row) or on YCB/BSA agar (bottome row) for 7 days at 35°C.

**Figure S4 (related to Figure 3).**
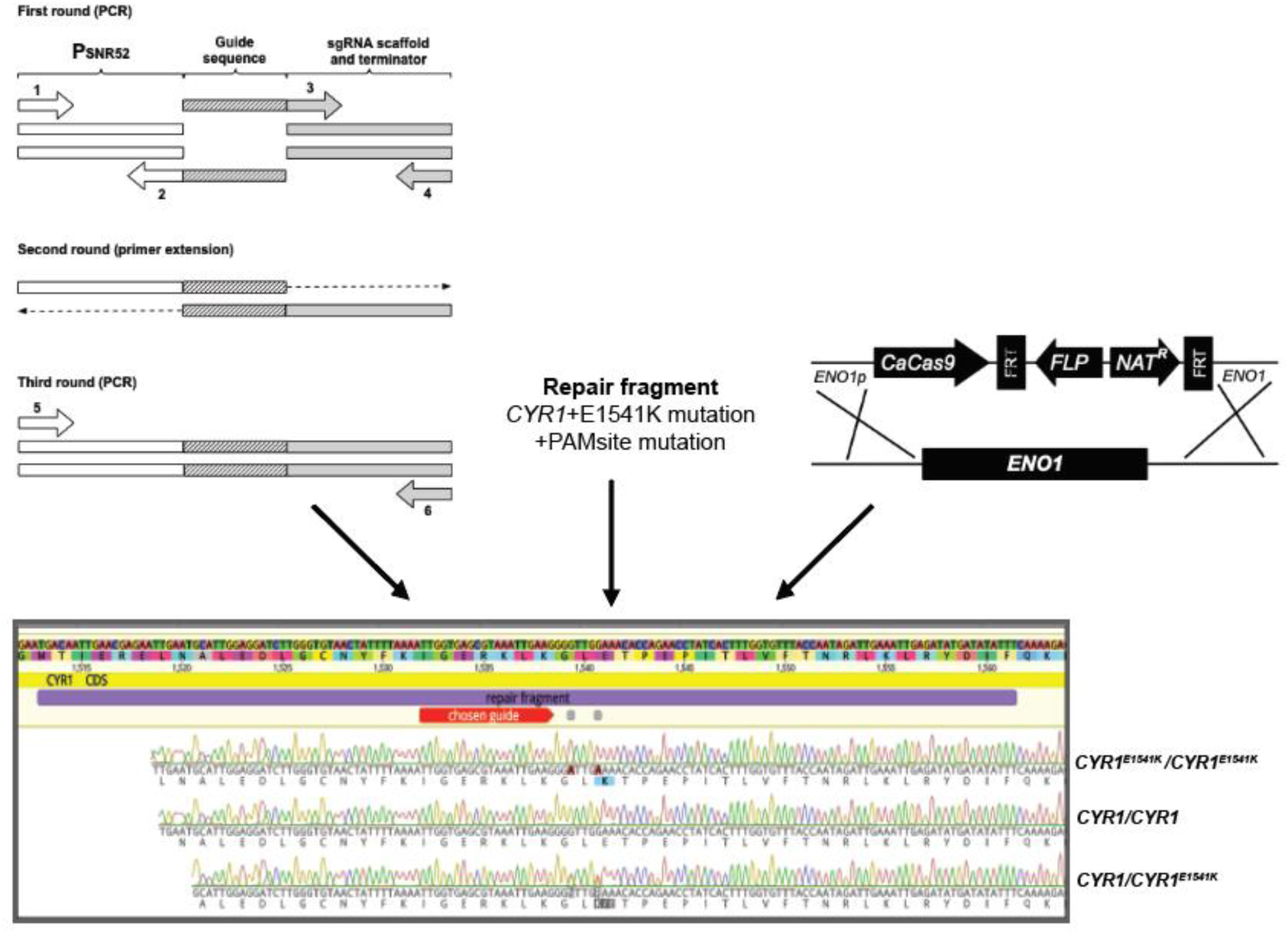
Strategy to introduce the *CYR1*^E1541K^ mutation in parental strain 101 using CRISP/Cas9. First, the guided RNA was produced followed by introduction of the repair fragment that harbors the *CYR1* single mutation. Then *C. albicans* strains were transformed by electroporation. Display of the sequence analysis demonstrated successful introduction of *CYR1* single mutation.

**Figure S5 (related to Figure 4).**
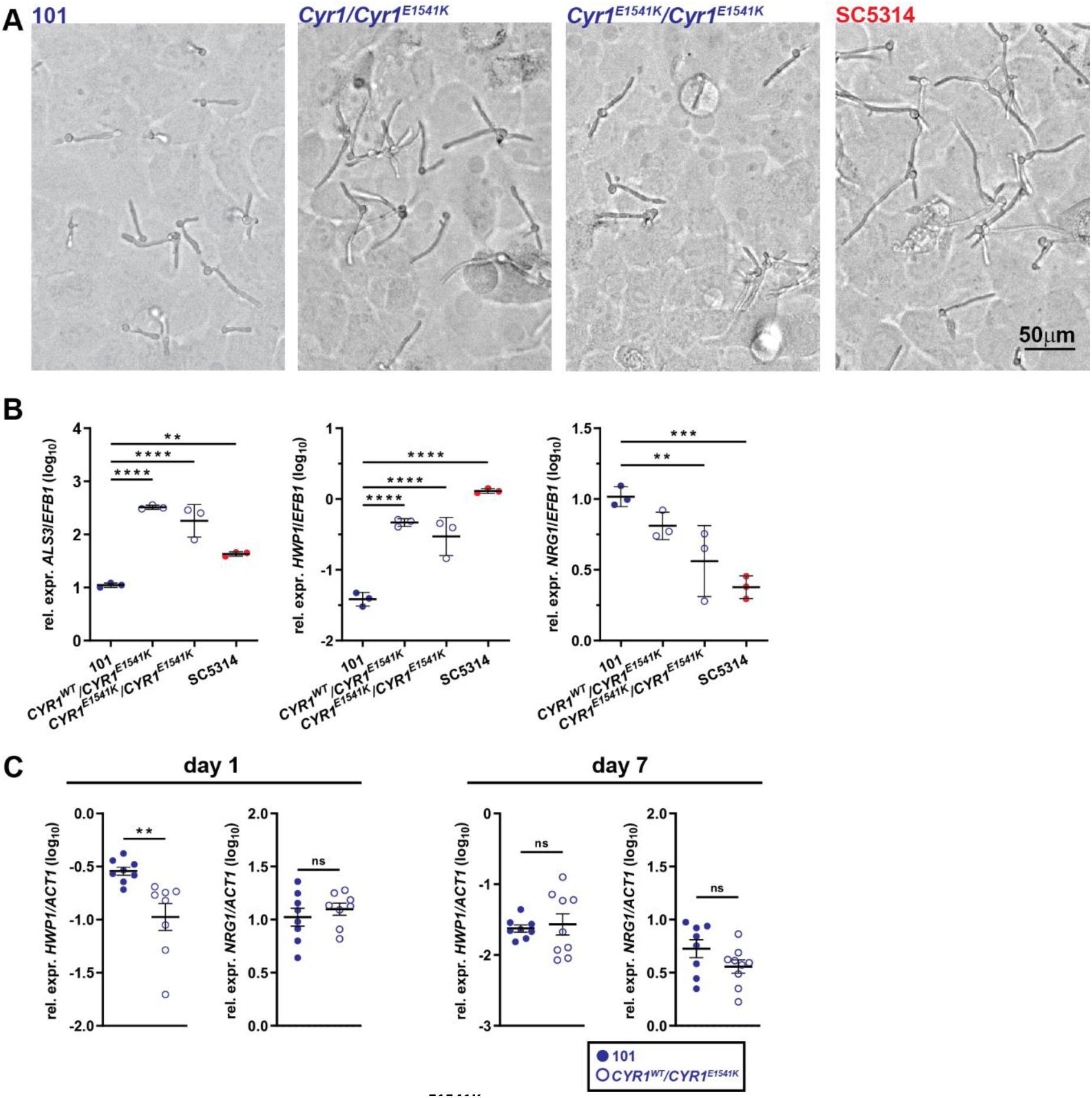
*CYR1*^E1541K^ induces the expression of virulence traits in the low-virulent strain 101. **A.** Representative images of filamentation *C. albicans* strains 101 wild type, 101_*CYR1*^E1541K^, and 101_*CYR1*^E1541K/E1541K^ and SC5314 wild type put in contact with a monolayer of TR146 keratinocytes for 3.5 – 4 hours at 37°C, 5% CO_2_. **B.** *HWP1*, *ALS3*, and *NRG1* expression levels by the fungal strains after exposure to TR146 keratinocytes for 24 hours at 37°C, 5% CO_2_. Each symbol represents one sample. Data are from one representative out of two independent experiments with three samples per strain each. The mean ± SD is indicated. **C.** *HWP1* and *NRG1* expression levels in the tongue tissue of C57BL/6 mice on day 1 after colonization with 101 wild type or 101_*CYR1*^E1541K^. Data are pooled from two independent experiments. The mean ± SEM is indicated. Statistical significance was determined using one-way ANOVA (B) or unpaired Student’s *t* test (C). *p<0.05, **p<0.01, ***p<0.001, ****p<0.0001

**Figure S6 (related to Figure 4).**
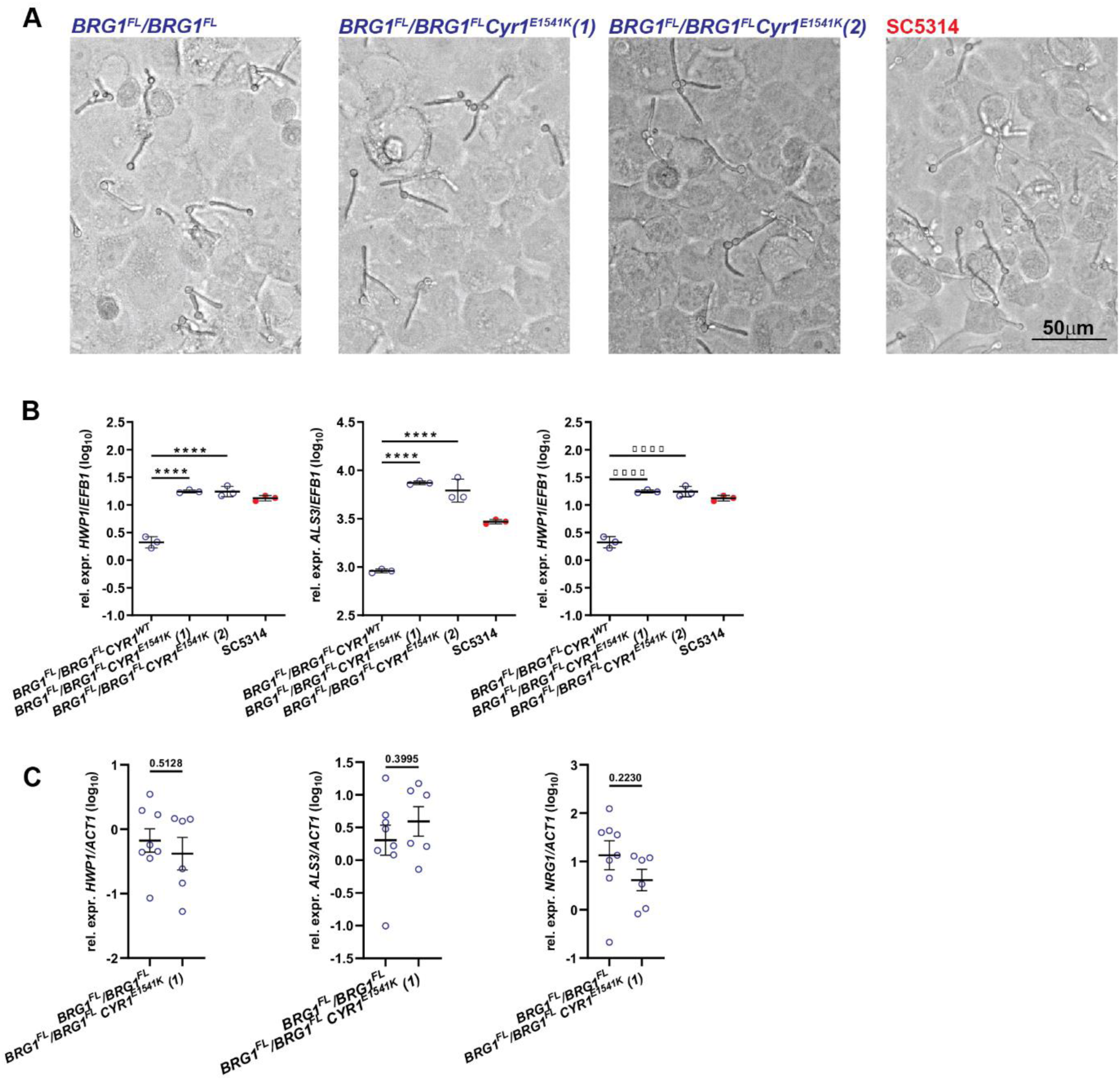
*CYR1*^E1541K^ -induced virulence traits are independent of the *BRG1* locus, and insufficient to elicit full virulence of strain 101 in the oral mucosa. A. Representative images of filamentation *C. albicans* strains 101_*BRG1*^FL/^*BRG1*^FL^, 101_*BRG1*^FL/^*BRG1*^FL^ *CYR1*^E1541K^ (clone 1 and clone 2), and SC5314 wild type upon contact with a monolayer of TR146 keratinocytes for 3.5 – 4 hours at 37°C, 5% CO_2_. . B. *HWP1*, *ALS3* and *NRG1* expression levels by the fungal strains after exposure to TR146 keratinocytes for 24 hours at 37°C, 5% CO_2_. Each symbol represents one sample. Data are from one representative out of two independent experiments with three samples per strain each. The mean ± SD is indicated. C. *HWP1*, *ALS3* and *NRG1* expression levels in the tongue tissue of C57BL/6 mice on day 7 after colonization with 101_*BRG1*^FL/^*BRG1*^FL^ or 101_*BRG1*^FL/^*BRG1*^FL^ *CYR1*^E1541K^ (clone 1). Each symbol represents one mouse. Data are pooled from two independent experiments. The mean ± SEM is indicated. Statistical significance was determined using one-way ANOVA (B) or unpaired Student’s *t* test (C). *p<0.05, **p<0.01, ***p<0.001, ****p<0.0001.

